# Diverse scientific benchmarks for implicit membrane energy functions

**DOI:** 10.1101/2020.06.23.168021

**Authors:** Rebecca F. Alford, Jeffrey J. Gray

## Abstract

Energy functions are fundamental to biomolecular modeling. Their success depends on robust physical formalisms, efficient optimization, and high-resolution data for training and validation. Over the past 20 years, progress in each area has advanced soluble protein energy functions. Yet, energy functions for membrane proteins lag behind due to sparse and low-quality data, leading to overfit tools. To overcome this challenge, we assembled a suite of 12 tests on independent datasets varying in size, diversity, and resolution. The tests probe an energy function’s ability to capture membrane protein orientation, stability, sequence, and structure. Here, we present the tests and use the *franklin2019* energy function to demonstrate them. We then present a vision for transforming these “small” datasets into “big data” that can be used for more sophisticated energy function optimization. The tests are available through the Rosetta Benchmark Server (https://benchmark.graylab.jhu.edu/) and GitHub (https://github.com/rfalford12/Implicit-Membrane-Energy-Function-Benchmark).

## Introduction

Accurate energy functions are critical for biomolecular structure prediction and design. Through physical, empirical, and statistical models of atomic and molecular interactions, these functions discriminate near-native from non-native conformations and optimize sequences to stabilize structures. Over the past 20 years, an influx of high quality structural data paired with new optimization tools have boosted the accuracy of soluble protein energy functions.^1^ A remaining task is to improve energy functions for membrane proteins, a class of molecules that constitutes 30% of all proteins^2^ and targets for over 50% of drugs.^3^

The heterogeneous lipid bilayer introduces several challenges for membrane protein energy function development. Biomolecular modeling tasks such as docking and design require benchmarking with both thermodynamic data and macromolecular structures. Yet, difficulties in over-expression and purification of membrane proteins have limited the quality and quantity of experimental validation data.^4^ For instance, membrane proteins represent less than 2% of structures in the Protein Data Bank^5^ and less than 1% of entries in ProTherm.^6^ Additionally, biomolecular modeling programs often accelerate calculations with implicit membrane models that represent the solvent as a continuous medium. This choice obfuscates the comparison between predictions and experimental data measured in a real lipid bilayer.

As a consequence, many membrane protein energy functions are trained for a single task on a small dataset. This strategy has been applied to various membrane protein modeling tools including estimating the ΔΔ*G* of mutation,^7^ hydrophobic thickness,^8^ native structure discrimination,^9,10^ refinement,^11^ protein design,^12^ symmetry,^13^ and protein-protein docking.^14,15^ These tools enabled a decade of membrane protein modeling; however, their generalizability is unclear. Small quantities of data prevent cross-validation: a technique that ensures performance on targets that are different from the training set. Further, small datasets may be feature-poor. For example, a set of membrane proteins may only contain transmembrane domains and exclude juxta-membrane domains important for function.

Recent advances have enabled energy function development with up to four training and testing sets. In the Rosetta community, this includes two new implicit membrane energy functions. The first model, developed by Weinstein *et al.*,^16^ was fit to transfer free energies from the dST*β*L assay^17^ and tested on datasets describing the folding and thermodynamics of single-span dimers. The second model is *franklin2019*, ^18^ our biologically realistic implicit membrane model that permits use of parameters for different lipid compositions. This model was fit with the Moon & Fleming hydrophobicity scale ^19^ and evaluated on four tests describing *α*-helical and *β*-barrel membrane proteins with complex topologies. Interestingly, Weinstein *et al.* reported excellent performance on homo-dimers of single transmembrane segments; however, the same model underperformed on our tests. This outcome demonstrates another important complication in this area: developers use different criteria to evaluate membrane protein energy functions, resulting in ambiguity.

For soluble proteins, these challenges are addressed by using multiple large, qualitatively and quantitatively diverse datasets that aim to fully describe the biomolecular system.^20^ For example, in the Rosetta soluble protein energy function, inclusion of both small-molecule thermodynamic data and macromolecule structures from X-Ray crystallography and NMR spectroscopy resulted in significant improvements to Lennard-Jones, electrostatic, and solvation parameters.^21^ The CHARMM force field is parameterized with a large collection of biophysical data.^22^ The Open Force Field project integrated multiple data sources with a Bayesian statistical framework to improve atom typing. ^23^ It would be ideal to apply a similar solution for membrane proteins. The closest examples involve parameterization of an anisotropic model for orientation prediction with free energy calculations^24^ and explicit lipid force fields for molecular dynamics with a combination of quantum mechanical calculations and experimental thermodynamic data.^25^ However, these examples do not yet incorporate benchmarks on macromolecule structures. Further, for membrane proteins there are only multiple small and non-diverse datasets that partially describe the system.

The goal of this work was to overcome this validation challenge by developing a set of sparse and diverse scientific benchmarks for evaluating membrane protein energy functions. We created tests that probe four areas of the membrane protein energy landscape: (1) protein orientation in the bilayer, (2) stability, (3) sequence, and (4) complete structures. The tests rely on a mixture of datasets that range in both size and quality, resulting in overall feature-rich optimization targets. Importantly, the tests are fast to evaluate to enable multiple iterations for optimization protocols. As a demonstration, we applied the scientific benchmarks to evaluate the accuracy of the *franklin2019* energy function. The tests identified energy function strengths and imperative areas for optimization. These results lay the groundwork for future energy function development and enable use of more sophisticated optimization tools such as deep learning.

## Results

To evaluate implicit membrane energy functions, we developed a set of 12 scientific benchmark tests (Table 1) In the following sections, we present each test with its dataset and we demonstrate the analysis with *franklin2019*, a current Rosetta implicit membrane energy function.

**Table 1:**
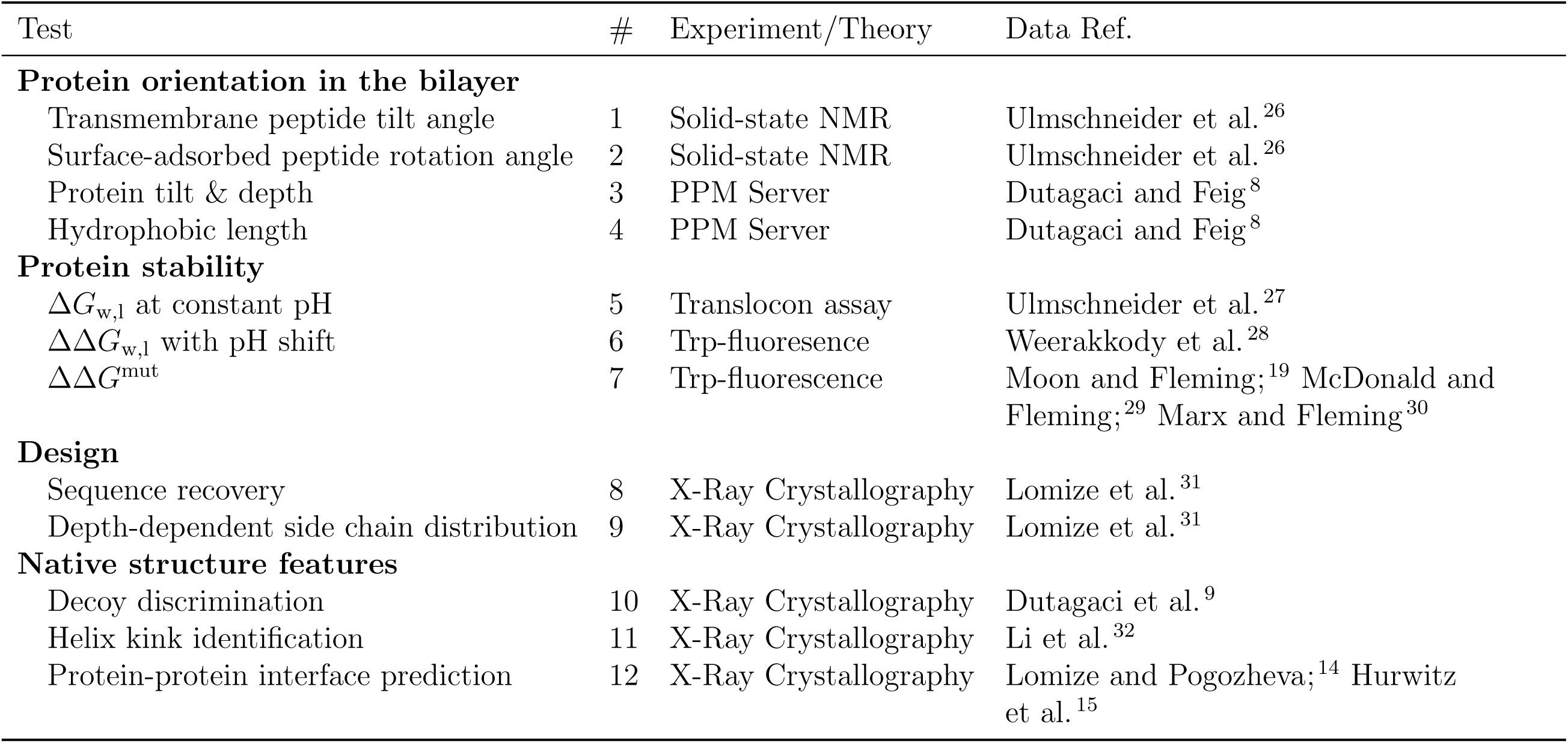
Membrane protein energy function benchmarks

### Test #1: Orientation of transmembrane peptides

Membrane proteins occupy precise orientations in the lipid bilayer to perform their biological functions. Thus, a key challenge for energy functions and the focus for our first tests is to recapitulate this position during structure prediction and design. This first test verifies that the most stable computed orientation of a transmembrane peptide corresponds to the native orientation. This test is the cornerstone of our benchmark because it was used for validation of early implicit membrane models.^33,34^ Here, the native orientation is defined as the tilt angle measured by solid-state NMR spectroscopy in the context of oriented lipid bilayers.^35^ To predict orientation, we developed a protocol to sample all possible orientations of the peptide relative to the implicit membrane within ± 60 Å of the bilayer center and tilt angles between ±180°(see Supplementary Information). The global energy minimum of all sampled positions is defined as the most stable predicted orientation for comparison to the experimental measurements. More generally, based on our biophysical intuition and the observation that transmembrane peptides typically prefer tilt angles between 0-45°, we expect those tilt angles and depths that span the membrane to be lower in energy relative to the aqueous phase or interface.

#### Dataset

The test set contains seven peptides with a single transmembrane domain (Table S2). The first four peptides are segments of biological membrane proteins from Ulmschneider et al. ^26^ The fifth peptide is the designed WALP23 peptide.^36^ Their tilt angles were measured in different lipid compositions including DPC micelles,^37^ DMPC vesicles,^38^ mixed DOPC:DOPG bilayers,^39^ and pure DOPC bilayers.^36^ While the experimental uncertainty is not available for all measurements,^26^ tilt angle measurements have a typical error range of ± 3-5°. In addition, to evaluate the preference of aromatic side chains for the membrane interface, we added to the set two designed poly-alanine helices with flanking tryptophan and tyrosine residues.^40,41^ Orientations have not been measured for these peptides, but predictions can still be compared to biophysical intuition.

#### Demonstration & Assessment

Test results with *franklin2019* are shown for a biological and a designed transmembrane peptide in Fig. 1a and 1b, respectively. The remaining results are shown in Fig. S1 and S2. We rated performance according to how many predictions fall within ± 10° of the measured value, a little more than the usual experimental error since some cases do not have reported errors.

**Figure 1:**
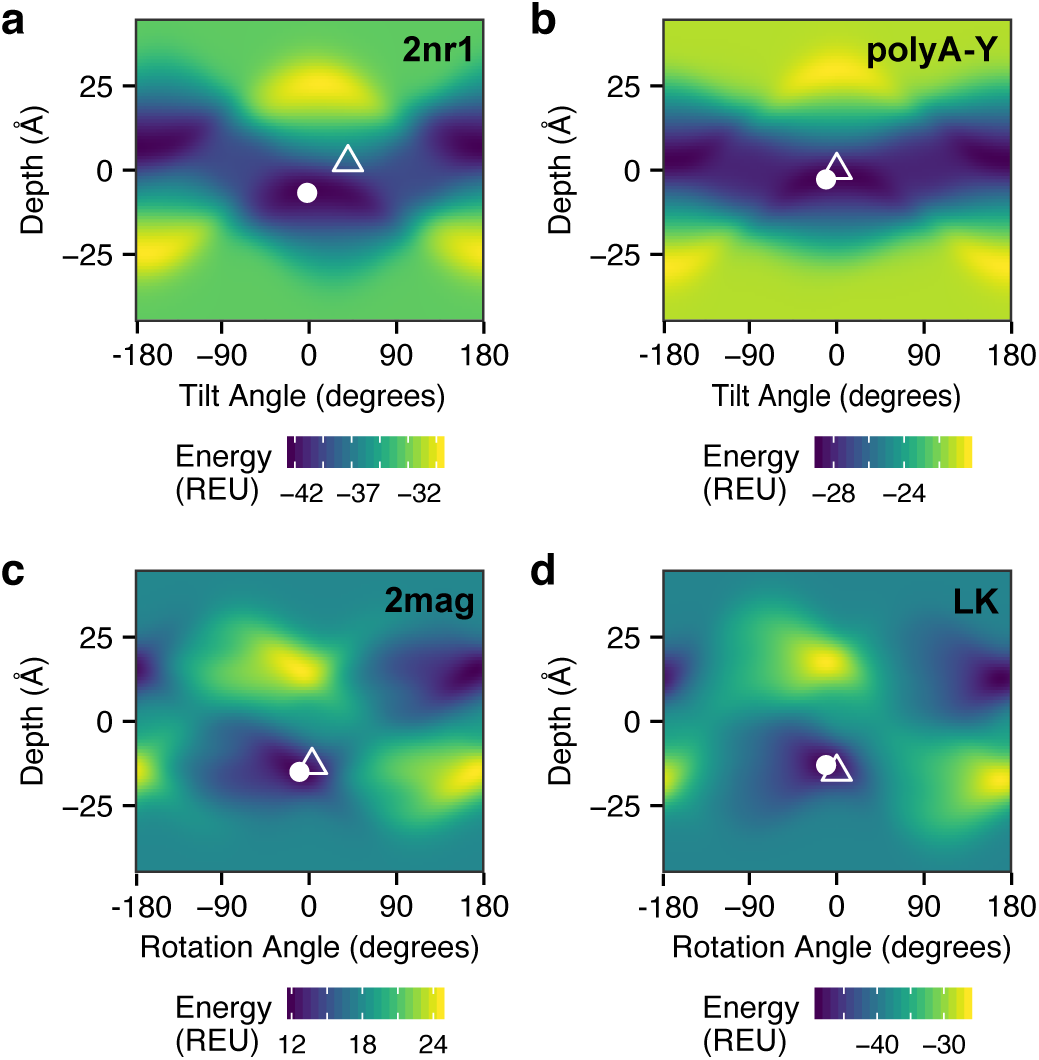
Implicit potentials capture the orientation of membrane associated-peptides with biological and designed sequences. A mapping between all sampled orientations and energies for (a) nicotinic acetylcholine receptor segment (b) poly-alanine trp-flanked peptide, (c) magainin, and (d) leucine-lysine repeat peptide. Tilt or rotation angle is on the *x*-axis, and depth relative to the bilayer center is on the *y*-axis. Each grid point represents a 1 Å and 1° and is colored by *franklin2019* energy, with low (favorable) energies in dark blue and high energies in yellow. The predicted lowest energy orientation is shown as a white circle, and where applicable the experimentally measured orientation is shown as an open triangle. Experimental measurements were only available for tilt angle and not depth; thus, we chose a depth of 0Åto represent this data.

As previously reported (Fig. 1 in Alford *et al.*^18^), *franklin2019* predicts the tilt angles of WALP and three of four biological peptides within 10°. For the poly-alanine aromatic-flanked peptides, the minimum energy occurs at a low energy tilt angle between 10-20°, which would reasonably expose the aromatic side chains to the aqueous solvent.

### Test #2: Orientation of membrane surface adsorbed peptides

Many proteins perform their functions by binding to the membrane surface at a specific orientation. This test verifies that the most stable surface-adsorbed peptide orientation corresponds to the native orientation. We define the native orientation by the rotation of the peptide relative to the helix axis and a specific membrane depth. Similar to **Test #1**, we sampled all pairs of membrane depths and rotation angles (SI Appendix).

#### Dataset

The dataset includes seven biological peptides^26^ and the designed leucine-lysine (LK) repeat peptide^42^ (Table S3). The experimental reference data are rotation angles measured by solution NMR in dodecylphosphocholine micelles or trifluoroethanol.^43^ In the experiments, rotation angles are reported with an uncertainty of ± 6-12°.^26^

#### Demonstration & Assessment

The mapping of orientations to energies for magainin and the LK repeat peptide calculated in an assumed DLPC bilayer are shown in Fig. 1c and Fig. 1d, respectively. The remaining targets are shown in Fig. S3. We rated performance according to how many predictions fall within the maximum experimental uncertainty across all of the measurements, i.e., 12°.

Fig. 1c-d and Fig. S3 show the energy landscapes with the predicted low-energy orientation (white circle) overlaying the measured value (white triangle). The landscapes reveal a repeating pattern of high energy (yellow) and low energy patches (dark blue) near the membrane interfaces (±18-23 Å). This pattern matches the biophysical intuition that the non-polar side of amphipathic peptides is more compatible with the membrane surface than aqueous solvent. Where experimentally measured values are available, the *franklin2019* energy function calculated rotation angles within ±12° of the native value for all of the surface-adsorbed peptides where experimentally measured values were available.

### Test #3: Orientation of multi-pass *α*-helical and *β*-barrel proteins

Accurately predicting peptide tilt angle is an important step toward predicting membrane protein orientation; yet, most membrane proteins have multiple transmembrane segments. Here, we examine how implicit membrane energy functions recapitulate the orientations of *α*-helical and *β*-barrel protein domains with complex topologies. We sample protein orientations (Fig. 2a-c) using the protocol described in **Test #1**. Then, because this data set is larger, we summarize the difference between the reference and calculated values across the set by computing the cumulative distribution of residuals.

**Figure 2:**
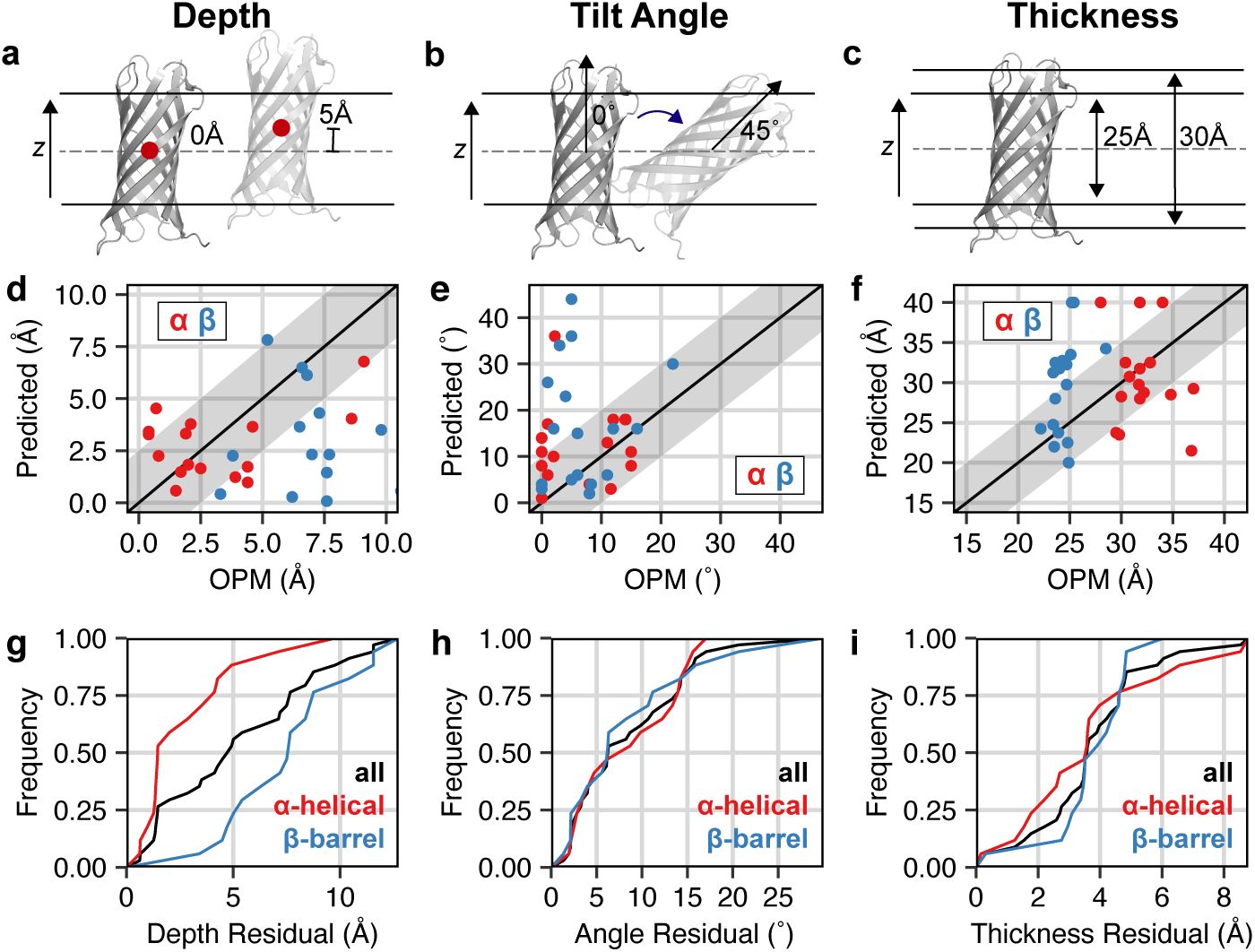
Protein orientation predictions of *α*-helical and *β*-barrel proteins. Comparison of membrane protein depth (a), tilt angle (b), and hydrophobic length (c) predicted by *franklin2019* relative to values predicted by the OPM anisotropic model.^31^ A *y* = *x* line is shown in each plot, with a light gray stripe for an error range of 2.5Å, 10° and 5 Å respectively. Targets that are *α*-helical (*β*-barrels) are represented as red (blue) points. Cumulative distribution for residuals of (d) membrane protein depth, (e) tilt angle, and (f) hydrophobic thickness. A distribution for all proteins is shown in black, and the distributions for *α*-helical only and *β*-barrel only are shown in red and blue respectively. Points at 40 Å indicate that full burial of the protein in the bilayer minimizes the energy.

#### Dataset

This test set was curated by Dutagaci and Feig ^8^ and includes 18 *α*-helical and 17 *β*-barrel proteins (Table S5). In the dataset, 60% of proteins reside in the *E. coli* outer membrane and 14% reside in the eukaryotic plasma membrane. The remaining 26% reside in either the mitochondria inner membrane, archea, or the *E. coli* inner membrane.

The most challenging step of this test is choosing reference data. There are various methods for identifying protein tilt angles and penetration depths in lipid bilayers or detergents including chemical modification, spin labeling, NMR, X-Ray scattering, and electron cryomicroscopy.^44^ The data vary in quality and have different assumptions and error sources. For this reason, the current reference data source is the Orientations of Proteins in Membranes database of membrane protein structures positioned in a 1,2-dioleoyl-sn-glycero-3-phosphocholine (DOPC) bilayer (OPM^31^). OPM computes the spatial position of a membrane protein using PPM; a method that combines an all-atom representation of the protein, an implicit anisotropic model of the lipid bilayer, and a universal solvation model. The final orientation is determined by minimizing the water-to-bilayer transfer energy, which is computed as a sum of van der Waals, electrostatics, and hydrogen bonding energies. ^24^ Using uncertainties from multiple original experiments, the estimated accuracy of OPM reference data is ±10° for tilt angle and ±2.5 Å for depth.^44^

#### Demonstration & Assessment

Membrane protein orientations predicted by *franklin2019* in a DOPC bilayer are given in Fig. 2d-e and the cumulative distribution of residuals is shown in Fig. 2g-h. We rated performance by the number of predictions that fall within the maximum uncertainty ranges of ±10° for tilt angle and ±2.5 Å for depth. For *α*-helical proteins, protein depth was correctly predicted for 62.5% of targets. In contrast, predictions were correct for 4% of *β*-barrel protein targets. Tilt angle prediction was more consistent, with 60% and 70% of protein tilt angles predicted within 10° of the OPM value for *α* and *β* proteins, respectively. Still, for *β*-barrel proteins, *franklin2019* predicts several outliers. For example, the predicted tilt angle for the alginate export protein (AlgE, 4afk) was 34°, whereas the OPM predicted value is 3°. The *franklin2019* prediction is likely unrealistic as it buries the pore entry. Overall, the data reveal that *franklin2019* predicts membrane depth more accurately for *α*-helical proteins than *β*-barrel proteins.

### Test #4: Membrane protein hydrophobic thickness

To overcome unfavorable exposure of non-polar side chains to water, membrane proteins generally have a hydrophobic thickness compatible with the bilayer thickness. Thus, hydrophobic thickness is an important parameter for predicting orientation and stability. In this test, we predict hydrophobic thickness by placing the protein at its OPM orientation and then iteratively rescoring the protein at membrane thickness values ranging from 10-40 Å (Fig. S4). The protein hydrophobic thickness is then defined as the bilayer thickness that minimizes the energy. This test uses parameters for a phosphatidylcholine head group.

#### Dataset

This test uses the same proteins as **Test #3**. Experimentally, hydrophobic thickness is measured using various techniques that often introduce different assumptions and errors. For instance, the experimentally measured hydrophobic thickness of OmpX (1qj8) was determined assuming that the protein does not tilt.^45^ For this reason, we use predicted hydrophobic thickness values from the OPM database^24^ as our reference. OPM reports thickness as the bilayer thickness that minimizes the water-to-bilayer transfer energy. The estimated uncertainty for hydrophobic thickness values from OPM is ± 2.5 Å.^44^

#### Demonstration & Assessment

A comparison of *franklin2019* and OPM is shown in Fig. 2f and Fig. 2i. The hydrophobic thickness was within 2.5 Å of OPM for only 30% of *α*-helical proteins and 10% of *β*-barrel proteins. At 5 Å, both categories improve to 75%; however, this threshold is large relative to the thickness differences between varied lipid compositions. For 5 targets, we observed that the energy continued to improve up to the limit of 40 Å. These points are plotted at T = 40 Å and indicate that *franklin2019* prefers to completely bury these proteins in the membrane. Overall, prediction of hydrophobic thickness was less reliable than orientation.

To better understand areas for improvement, we examined the outliers in Fig. 2f. The most incorrect prediction is for the bacterial semiSweet transporter (4×5n, Fig. 2f). Here, *franklin2019* predicted a hydrophobic thickness value of 21.5 Å whereas OPM predicted a value of 36.8 Å. This suggests error in pore estimation because *franklin*2019 leaves the entry and exit of the pore more exposed than OPM.^31^ The five targets for which no minimum energy hydrophobic thickness was found are photosynthetic reaction center (1rzh), potassium channel KcsA (1r3j), opioid delta receptor (4n6h), outer membrane protein A (1qjp), and the hemoglobin binding protease autotransporter (3aeh). It is less clear why these cases were outliers. Both the photosynthetic reaction center and opioid delta receptor have significant juxta-membrane domains, suggesting that *franklin2019* ‘s implicit membrane interfacial representation may be insufficient to differentiate transmembrane from non-transmembrane segments. Another possibility is that pore estimation further complicates predictions.

### Test #5: Stability of transmembrane peptides at neutral pH

The next three tests focus on recapitulating thermodynamic properties of membrane proteins. In cells, the translocon machinery is responsible for recognizing transmembrane segments and integrating them into the bilayer. Once folded, membrane proteins remain in the bilayer due to a favorable water-to-bilayer transfer free energy. To keep the protein in the membrane, we therefore must accurately estimate the transfer energy from solvent to the bilayer. In **Test #5**, we build on the calculations sampling all possible peptide orientations from **Test # 1**. Then, we compute the Δ*G* of insertion as the energy difference between the lowest energy orientation of the peptide in the lipid bilayer phase and in the aqueous phase (see Supplementary Information) to compare to experimental insertion energy measurements.

#### Dataset

The test set includes five poly-leucine peptides designed by Ulmschneider et al. ^27^ There are four peptides in the set that follow a *GL*_*X*_*RL*_*X*_*G* motif, where *X* = 5, 6, 7, 8 (Table S6). The fifth peptide follows a different motif pattern by adding a flanking tryptophan: *GWL*_6_*RL*_7_*G*. The reference transfer energies are taken from molecular dynamics simulations in POPC bilayers that were validated against intrinsic fluorescence measurements. The experimental uncertainty of the measured Δ*G*_ins_ values is ±1.4 kcal mol^−1^.

#### Demonstration & Assessment

The *franklin2019* energy function computes the water-to-bilayer insertion energy using a Gibbs insertion energy approximation. As in many other Rosetta based calculations, we expect this energy function may reproduce trends but not exact values. Thus, we evaluate the ability to reproduce trends using the Pearson correlation coefficient (*R*^2^) between the experimentally measured and predicted values (although as accuracy improves in the future, exact predictions can also be evaluated). In calculating *R*^2^, we use a Grubbs test to eliminate outliers.

A comparison between the *franklin2019* prediction in POPC bilayers and reference Δ*G* values from molecular dynamics (MD) is shown in Fig. 3a. Encouragingly, the Pearson correlation coefficient is high (*R*^2^ = 0.996), and no points are outliers. The slope of the best fit line is high (2.41 REU-mol/kcal), revealing that while the relative energies are correct, *franklin2019* overestimates the overall benefit of insertion. Further, the comparison indicates a reference state calibration issue because the Δ*G* of inserting the GL5 and GL6 peptides is favorable; whereas, the MD indicates insertion is unfavorable.

**Figure 3:**
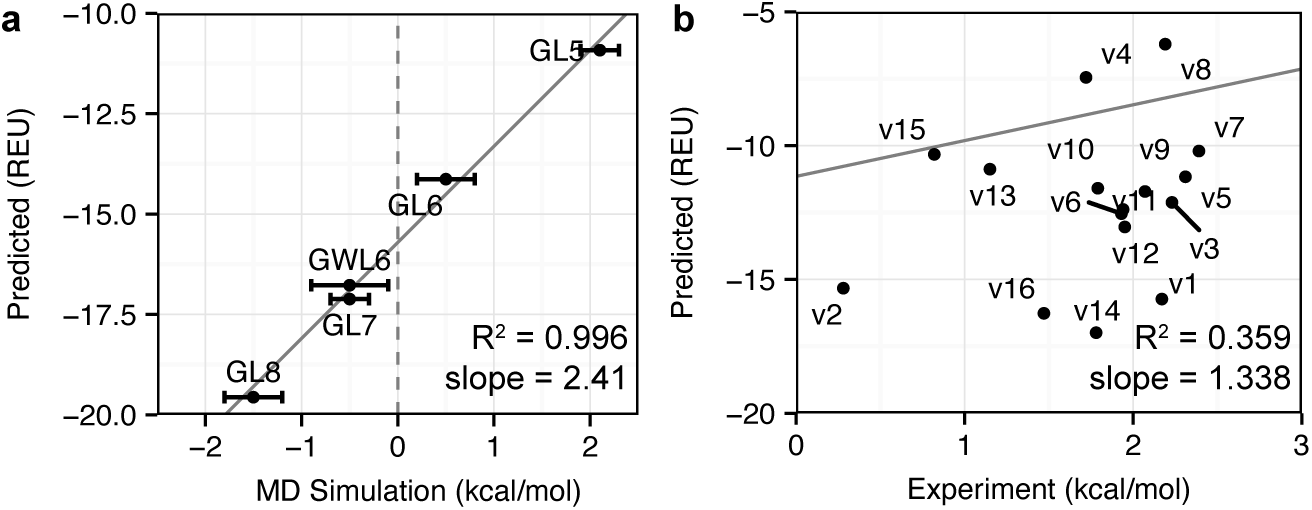
Comparison of predicted and experimentally measured peptide insertion energies. (a) Prediction of the ΔΔ*G* of insertion for five designed poly-leucine peptides of varying length and flanking residues from Ulmschneider et al. ^27^ (b) Prediction of the ΔΔ*G* of insertion upon pH shift from pH 8 to 4 for 16 variants of the pH-sensitive low insertion peptide sequence from Weerakkody et al. ^28^ Experimental measurements were taken by intrinsic fluorescence in POPC liposomes and calculations were performed for DLPC.

### Test #6: Stability of transmembrane peptides at acidic pH

The water-to-bilayer transfer energy is influenced by many factors including changing conditions in the intracellular and extracellular environment. Physiologically, extracellular acidity is an important biomarker for tumor growth and development.^46^ To benchmark the ability of the energy function to capture pH, we evaluated the prediction of peptide insertion energy when there is a change in pH. The test performs two grid-style searches using the protocol described in **Test #1** with Rosetta-pH^47^ to find the best peptide orientation at both pH 4 and 8 (see Supplementary Material). Then, the insertion energy difference is computed as the difference between the peptide integrated into the bilayer at pH 4 and in the aqueous phase at pH 8 (i.e., ΔΔ*G*_*p*H,ins_ = Δ*G*_ins,*p*H4_ − Δ*G*_ins,*p*H8_, see Supplementary Material).

#### Dataset

The test set includes seventeen peptides designed by Weerakkody et al. ^28^ that insert into membranes upon a shift from neutral to acidic pH, called pHLIP peptides (Table S7). The reference value is the water-to-bilayer transfer energy measured using titration and fluorescence experiments in the context of POPC liposomes. The experimental uncertainty of the measured ΔΔ*G*_pH,ins_ values is ±1.0 kcal mol^−1^.

#### Demonstration & Assessment

A comparison of the *franklin2019* and experimentally measured values is given in Fig. 3b. Similar to **Test #5**, we aim to maximize the Pearson correlation coefficient. In contrast to **Test #5** at neutral pH, the correlation between the experiment and calculation is poor, and the energy function prefers the peptides in solution rather than in the bilayer. We suspect two sources of problems. First, the underlying pKa values do not account for the membrane. Second, the *franklin2019* Coulomb term does not account for changes in the dielectric constant in the membrane. This is a critical area of future energy function optimization because the shifted pK_*a*_ values in the bilayer affect the stability of membrane proteins at all pH values.

### Test #7: ΔΔ*G* of mutation

Test #7 evaluates how changes to the sequence of a membrane protein affect its overall thermostability. This quantity, called ΔΔ*G*_mut_, is a critical building block for membrane protein design and evaluating the effects of genetic mutations on protein function. To predict ΔΔ*G*_mut_, we used our previously described fixed-backbone and fixed-orientation protocol^48^ that evaluates the difference in total energy between the mutant and wild-type (see Methods).

#### Dataset

We used three sets of ΔΔ*G* measurements from the Fleming lab. All of the measurements were taken at equilibrium in DLPC vesicles and in the context of a *β*-barrel protein scaffold. The three datasets are: (1) mutations from alanine to all 19 remaining canonical amino acids at a lipid-exposed site on OmpLA^19^ (2) mutations from alanine to all remaining 19 canonical amino acids at a lipid-exposed site on PagP^30^ and (3) mutations from alanine to tryptophan, tyrosine, or phenaylanine at different membrane depths on OmpLA.^29^ The experimental uncertainty of the measured ΔΔ*G*_mut_ values is ±0.6 kcal mol^−1^.

#### Demonstration & Assessment

We previously reported the performance of *franklin2019* on the OmpLA and PagP datasets.^18^ Here, we focus on the third set which probes the contribution of aromatics to stability. ^40,41^ To emulate the experiment we modeled an implicit DLPC bilayer. A comparison of the *franklin2019* and experimentally measured values is shown in Fig. 4, focusing on tryptophan and tyrosine because both demonstrate strong depth-dependence.^29^ Unfortunately, there was no correlation between the predicted and experimentally measured values. The Pearson correlation coefficient was *R*^2^ = −0.135 for tyrosine and *R*^2^ = 0.023 for tryptophan.

**Figure 4:**
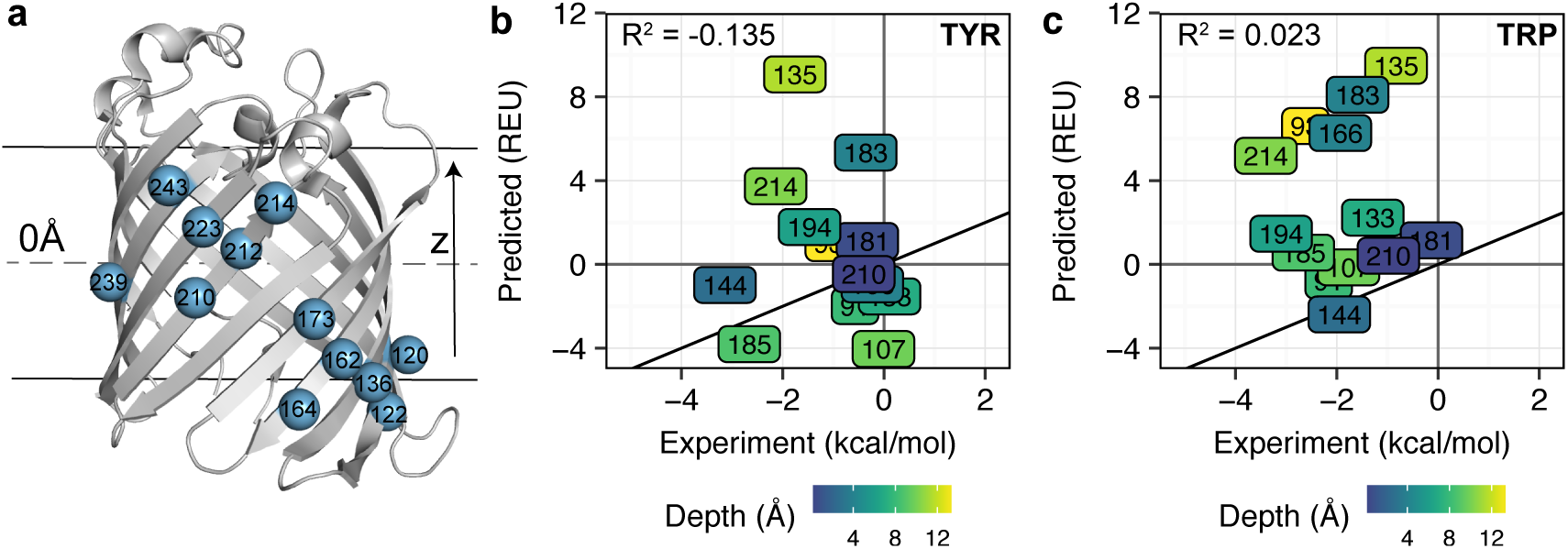
Comparison of predicted and experimentally measured depth-dependent ΔΔ*G*_mut_ values. (a) Outer membrane protein phospholipase A (OmpLA, PDB 1qd6) with each host site highlighted in blue. A comparison of predicted and experimentally measured ΔΔ*G*_mut_ for mutations to tyrosine and tryptophan are shown in (b) and (c) respectively. Each position is colored by depth relative to the membrane center plane in Å on a scale from blue (closer to the center) to yellow (closer to the interface/water barrier). The *y* = *x* line is shown as a bold black line. The experimentally measured values were taken from McDonald and Fleming.^29^

Interestingly, the Pearson correlation coefficient between the measured values and the *franklin2019* water-to-bilayer energy term was higher than that to the full energy function, with *R*^2^ = 0.451 for tyrosine and *R*^2^ = 0.608 for tryptophan (Fig. S5). To explore whether the ΔΔ*G* was driven by factors other than membrane heterogeneity, we mapped the contribution of all component energy terms to the ΔΔ*G* value (Fig. S6 and Fig. S7 show all values over 0.01 REU). The largest contributions were from rotamer energies, suggesting that steric clashes between guest side chains and neighboring side chains inflated the cost of substitution.

### Test #8: Sequence Recovery

The next two tests concern the ability to identify residue types in design calculations. Test #8 probes whether the energy function can recover native membrane protein sequences. To run the test, we perform redesign using Rosetta’s Monte Carlo fixed-backbone design protocol.^20^ Each protein is initialized in the orientation from the OPM database,^31^ and the orientation remains fixed. For simplicity, we use a DLPC membrane in our calculations. We compute two metrics: (1) the fraction of amino acids recovered (sequence recovery) and (2) the divergence of the designed amino acid distribution from the native amino acid distribution (Kullback-Leibler divergence). To tailor this test for the membrane, we also compute these metrics for subsets of amino acids exposed to the aqueous phase (outside of membrane or pore facing), lipid phase, and interfacial region. An optimal energy function would maximize sequence recovery and minimize divergence.

#### Dataset

The test set includes 133 *α*-helical and *β*-barrel membrane proteins. The starting dataset was curated by Koehler Leman et al. ^49^ and revised to only include proteins with known subcellular localization. ^18^ In this set, all entries have resolution of 3.0 Å or better, and no two sequences share more than 25% sequence identity. The native and host lipid compositions for these proteins vary widely and include compositions not yet covered by the *franklin2019* lipid parameters. So here for simplicity, we perform all design calculations in a DLPC bilayer.

#### Demonstration & Assessment

We previously reported the performance of *franklin*2019 on the sequence recovery test (Fig. 4 in Alford et al.).^18^ Here, the recovery rate was high (31.8%) and the divergence between designed and native residue distributions was low (KL = −2.7). Both metrics improved relative to prior energy functions.

### Test #9: Depth-dependent side-chain distribution

Test #9 evaluates whether the energy function captures bilayer depth-dependent features of membrane protein sequences. This test has been used previously for calibrating statistical implicit membrane potentials.^16,17,50^ To run the test, we first generate redesigned proteins using the same protocol as in **Test #8**. We use a kernel-density estimate to compute the depth-dependent (*z*-dependent) distribution of amino acids in both the native and redesigned proteins. We then use numerical integration to compute the difference in the area under the curve (AUΔC) between the native and redesigned distributions. An effective implicit membrane energy function qualitatively matches the shape of the distribution and minimizes the AUΔC difference.

#### Dataset

This test uses the same set of 133 protein structures as in **Test #8**.

#### Demonstration & Assessment

The depth-dependent amino acid distributions for proteins redesigned with *franklin2019* are shown in Fig. 5, and the AUΔC values are shown in Fig. S8. The profiles reveal both native-like and non-native-like properties of *franklin2019*. The best predicted amino acid distributions were for polar, non-polar, and some aromatic amino acids, namely T, Ala, Pro, Phe, Tyr, Met, Ile, Asn, and Gly (Fig. S8; AUΔC *<* 0.01). In contrast, Leu and Val were over enriched in the membrane core, resulting in larger AUΔC values of 0.018 and 0.022, respectively. The disparities were larger for charged amino acids. Specifically, the presence of Asp and Glu in the membrane is underestimated, whereas the presence of Arg and Lys is overestimated. Similarly, lack of asymmetry in the distributions reveals that *franklin2019* does not capture the “positive-inside rule”, which dictates that cytosolic loops near the bilayer contain more positively charged amino acids.^17,51^ Further, the distribution misses enrichment of positively charged side chains in the inner leaflet. Both of these features have been observed using the Elazar energy function. ^17^

**Figure 5:**
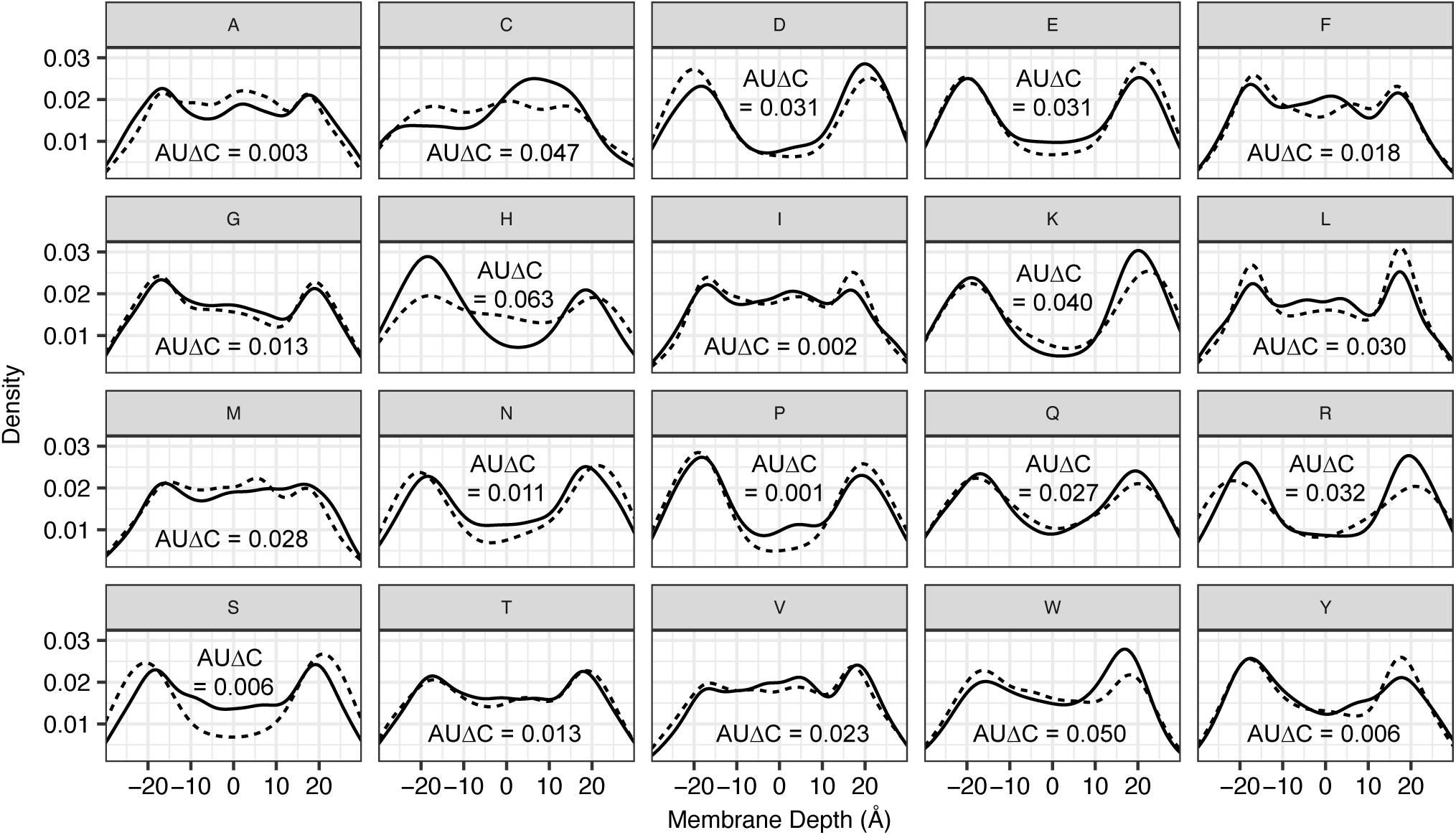
Comparison of the depth-dependent side chain distribution in native and designed protein sequences. The native amino acid distribution is shown with a solid line and the designed amino acid distribution is shown with a dotted line. Each panel represents the distribution for one of the twenty canonical amino acids with the membrane depth ranging between −30 Å and 30 Å, and the AUΔC value shown is the absolute area between the two distributions.

### Test #10: Native decoy discrimination

A key task for membrane protein energy functions is to distinguish near-native from non-native backbone structures. So, the last three tests involve discriminating between alternate structures. In previous work, there were two molecular dynamics discrimination studies with implicit membrane models.^9,10^ Recently, we also reported structure discrimination results for *franklin2019*.^18^ Here, we expand the test by increasing the number and structural diversity of decoy models. The native discrimination test is performed by refining a set of decoy models in the context of the candidate energy function. Then, we quantify discrimination using the Boltzmann-weighted average root-mean-squared-deviation (RMSD) over all models (*D*). We also qualitatively examine the ranking of decoys by score and RMSD. An optimal energy function would exhibit a low *D* and funnel-like arrangement of decoys, with non-native decoys assigned high energies and near-native decoys assigned low energies.

#### Dataset

The dataset includes five targets: bacteriorhodopsin (brd7), fumarate reductase (fmr5), lactose permease (ltpa), rhodopsin (rhod), and V-ATPase (vatp). The resolution of each crystal structure is 1.8 Å for brd7, 1.78 Å for fmr5, 3.5 Å for ltpa, 2.2 Å for rhod, and 2.1 Å for vatp. Each target is represented by decoys from two datasets: (1) the Dutagaci et al. set^9^ includes decoys between 4–14 Å RMSD from the native crystal structure, and (2) the Yarov-Yarovoy et al. set^52^ includes decoys between 5–40 Å RMSD.

To increase the number and diversity of decoys for each target, we used RosettaM-PRelax^48^ to generate five models from each decoy structure to provide 0.5–1.5 Å RMSD of additional variation. Since all of the X-Ray crystal structures were determined in detergents, we performed the calculations in a default lipid composition of DLPC. This bilayer is appropriate because all of the targets were expressed in *E. coli*.

#### Demonstration & Assessment

Fig. 6 summarizes the *franklin2019* native structure discrimination results for all five targets, and Table 2 lists the discrimination score for all decoys (*D*_all_), Dutagaci decoys (*D*_Dut_), and Yarov-Yarovoy decoys (*D*_YY_). Consistent with previous results,^18^ discrimination of Dutagaci decoys for each target is high and the models form a funnel. Interestingly, structure discrimination worsens with the addition of low-resolution decoys for two targets: bacteriorhodopsin and fumarate reductase. In both cases, there are models near 15 Å that score the same or better than models near 4 Å, suggesting that the energy function requires improvements to recognize native-like helix-helix contacts when there are large differences between possible conformations.

**Table 2:** Native structure discrimination by *franklin2019. D* is the Boltzman-weighted average RMSD over all models, with lower values indicating better identification of near-native models.

| Target | $D_{\text{all}}$ (Å) | $D_{\text{Dut}}$ (Å) | $D_{\text{YY}}$ (Å) |
| --- | --- | --- | --- |
| brd7 | 7.8 | 4.6 | 7.8 |
| fmr5 | 14.2 | 5.5 | 14.2 |
| ltpa | 4.8 | 4.8 | 13.7 |
| rhod | 4.3 | 4.3 | 12.7 |
| vatp | 4.0 | 4.3 | 4.0 |

**Figure 6:**
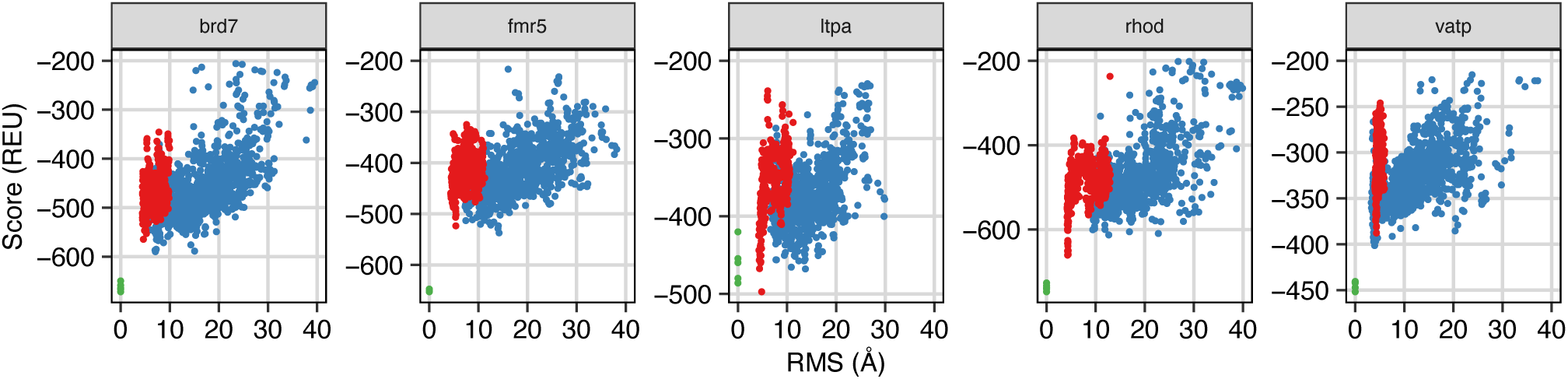
Decoy discrimination of five targets by the *franklin2019* implicit membrane potential. Each panel shows the decoys for each target ranked by energy (in REU) and RMSD (Å) of the C*α* atoms to the native (x-ray) structure. The high resolution (Dut) decoys (1–11 Å) are shown in red, the low resolution (YY) decoys (5–40 Å) are shown in blue, and the refined native structures are shown in green.

### Test #11: Helix kinks

A unique feature of membrane proteins is the distortion of *α*-helices into kinked and curved conformations. ^53^ Upon first look, kinked transmembrane helices seem counterintuitive because backbone hydrogen bonds are more stable in the membrane and kinks will break hydrogen bonds. In reality, there are multiple biochemical possibilities to resolve the hydrogen bonds, including using a proline,^54^ a vestigial proline,^55^ and non-canonical backbone hydrogen bonding patterns.^56,57^ To evaluate the energy function’s capacity to identify native non-canonical helical conformations, we scored conformational ensembles of membrane proteins with at least two conformational states with known structures where one structure exhibited a kinked helix that straightens in the second structure. First, assuming the system is stabilized by harmonic potentials,^58^ we generated conformational ensembles using normal mode analysis. We used KinkFinder^55^ to compute the kink angle, and then, to test whether the energy function can discriminate the native conformation, we calculated the energy of each model in the conformational ensemble.

#### Dataset

The dataset includes three targets: (1) potassium channel KcsA, (2) adiponectin receptor 1, and (3) platelet activating receptor (Table S8). The experimental kink measurements are derived from the crystal structure. For these cases, the native bilayer was ambiguous, so for all calculations we chose a DLPC system.

#### Demonstration & Assessment

A comparison of energies and kink angles for each state are shown in Fig. 7. As an example, we discuss the conformation of transmembrane helix 2 (TM2) in the the potassium channel KcsA (Fig. 7a). In the channel’s closed state, TM2 is kinked (red), whereas in the open state, TM2 is slightly curved (blue). However, Fig. 7a shows that the conformations generated from the open state where the native angle is 40° cluster closer to 20° and these conformations have a significantly higher score than the straight helix conformations. indicating that Rosetta prefers to straighten TM2 in the open state, even though both states are realistic. This error also occurs for the second target, adiponectin receptor 1 (Fig. 7b). These results suggest that improvements are needed to accurately capture helix conformations.

**Figure 7:**
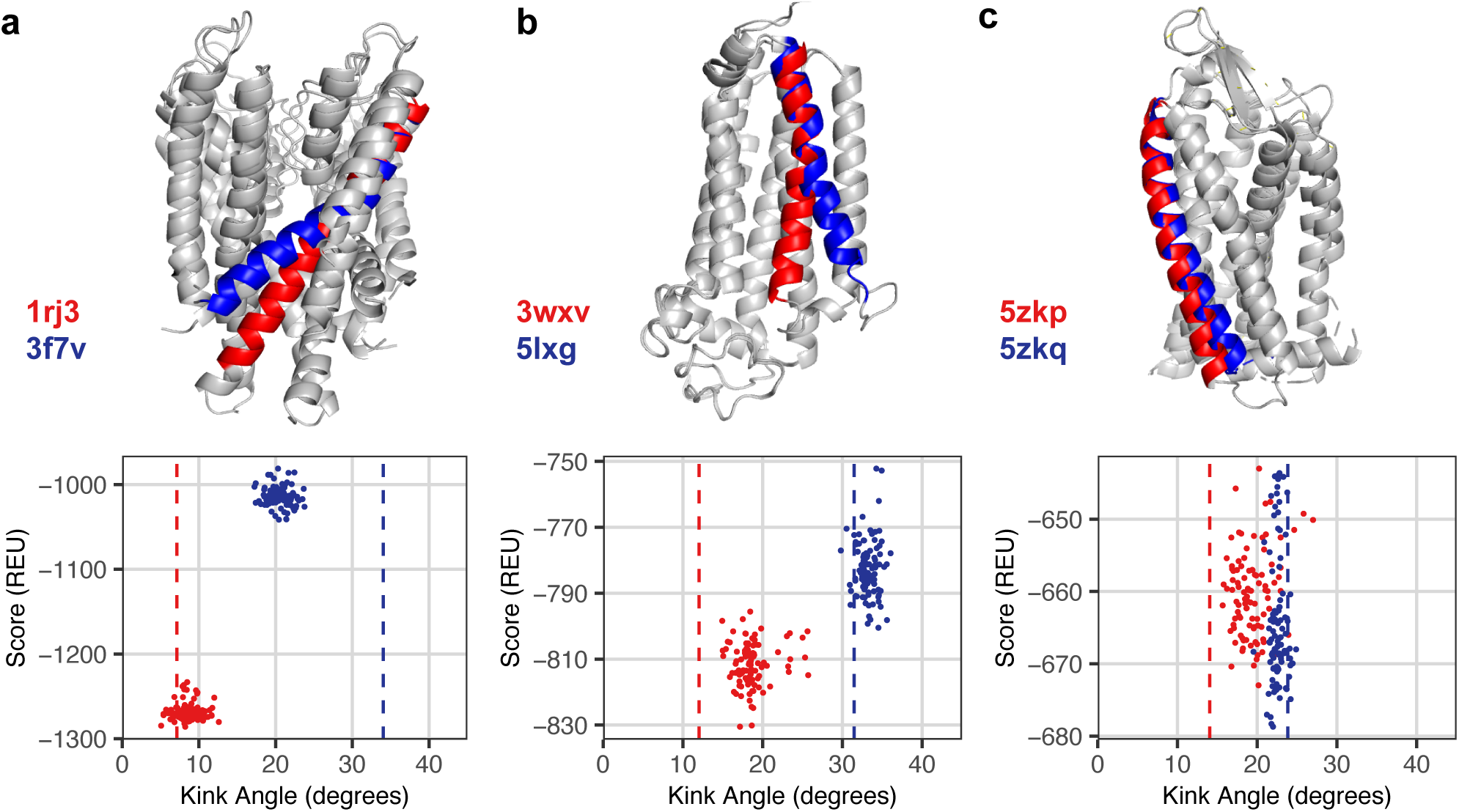
Kinked and straight conformations of *α* helices are not distinguished correctly. A structure discrimination experiment for kinked and straight helix conformations is shown for three targets: the open (3f7v) and closed (1r3j) conformation of the potassium channel KcsA (left), the active (3wxv) and inactive (5lxg) conformation of the adiponectin receptor, and (c) the active (5zkp) and inactive (5zkq) conformation of the platelet activating receptor. The top row shows the lowest energy conformation for both states and the bottom row ranks the energy of each conformation by kink angle relative to the native (denoted by a dotted line).

### Test #12: Membrane protein-protein interactions

A final challenging task for implicit membrane models is to distinguish near-native from non-native membrane protein-protein interface structures. A range of studies hint toward key interface features, including the GxxG motif,^59^ bifurcated *C*_*α*_ hydrogen bonds,^60^ and apolar side chain packing.^61^ However, there have only been a few general efforts to dock membrane proteins.^14,15,48,62^

Toward this goal, the final test is membrane protein-protein docking. We use RosettaM-PDock^48^ to generate low-energy orientation of the protein partners by performing rigid-body rotations and translations with cycles of side-chain repacking and torsion minimization (see Methods). For each target, we generated 5,000 candidate models and then used CAPRI metrics^63^ to evaluate the distance to crystal structures. We also computed two additional scoring metrics: (1) the number of near-native decoys in the 5 top-scoring decoys, ⟨*N* 5⟩, and (2) the enrichment of high quality models in the 1% and 10% top-scoring models,^64^ ⟨*E*_1%_⟩ and ⟨*E*_10%_⟩.^65^ Angle brackets denote bootstrapped averages over resampled decoy sets.

#### Dataset

This test uses three existing benchmark sets. The first dataset comprises 18 homodimers formed by single transmembrane helices.^14^ The second dataset comprises 48 homo- and hetero-dimers formed by multi-pass *α* helical proteins.^15^ The third dataset is a subset of the second consisting of nine targets with the starting backbone generated by homology modeling^66^ to simulate an unbound docking scenario where the starting conformations of the partner are not pre-configured in the bound state. All calculations were performed in a DLPC bilayer.

#### Demonstration & Assessment

Fig. 8 summarizes results for protein-protein docking with *franklin2019*. First, we examined the efficacy of docking structures from the crystalized bound state. We found that the membrane protein docking routine identifies high quality models for 90% of single-helix homodimer targets and 45% of multi-pass heterodimer targets, as indicated by two-fold enrichment in the top-scoring 1% of models (Fig. 8a-b). These targets are easy because the binding partners are already in the right conformation for binding. To evaluate interface recognition in more detail, we need to dock unbound targets.

**Figure 8:**
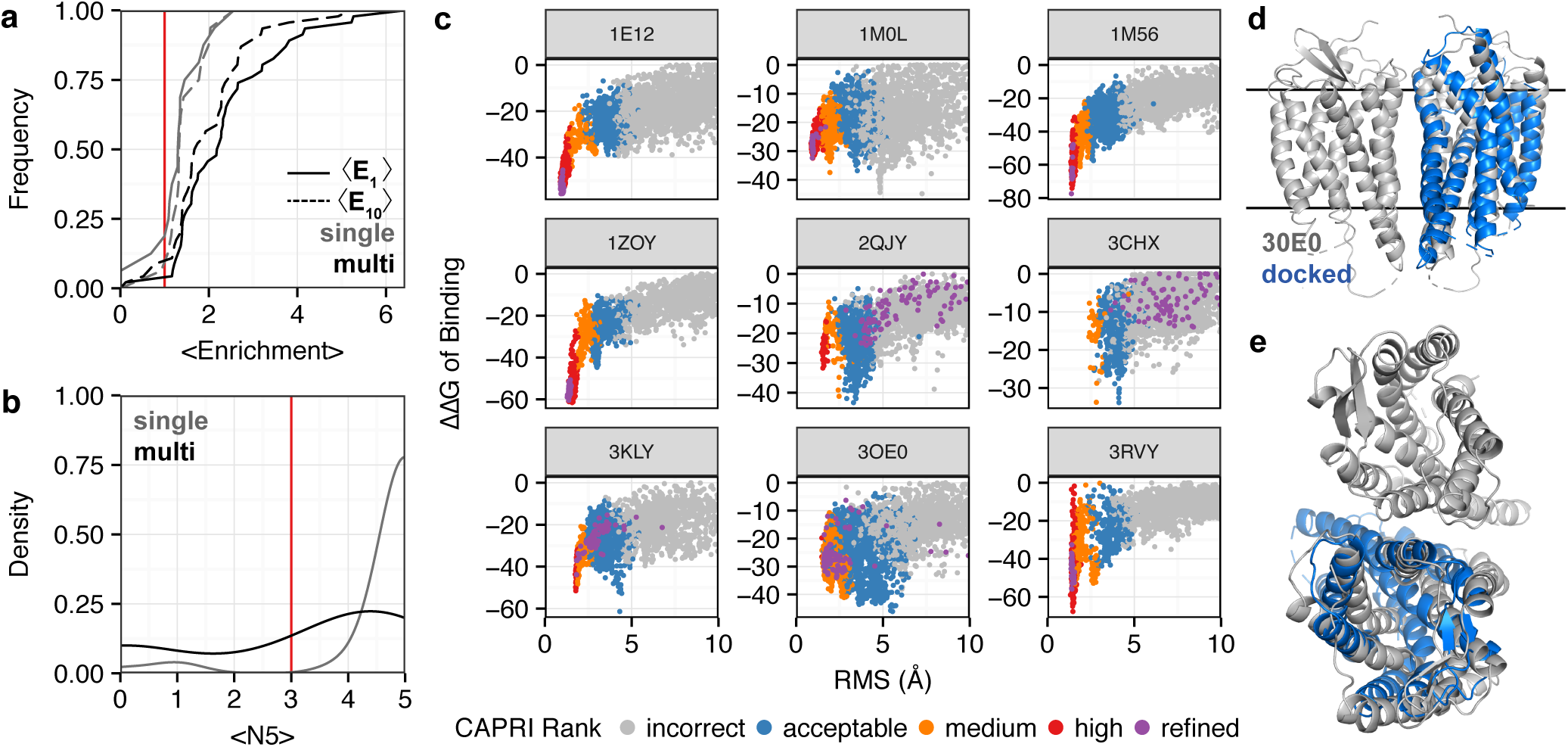
Implicit potentials identified native-like membrane protein-protein interfaces for nearly half of homology modeled targets. A summary of docking performance on bound (easy) targets is shown on the left hand side. (a) Cumulative distribution for the enrichment of high-quality models in the 1% and 10% top-scoring models, given as ⟨*E*_1%_⟩ (solid line) and ⟨*E*_10%_⟩ (dashed line), respectively. The distribution for single-transmembrane homodimers is shown in gray (n = 18) and the distribution for multi-pass homo- and heterodimers is shown in black (n = 48). (b) Cumulative distribution for the number of near-native decoys among the five top-scoring decoys, called ⟨*N*5⟩. Again, the distribution for single-transmembrane homodimers is shown in gray and the distribution for multi-pass targets is shown in black. The success cutoff of three models is shown in red. (c) Performance of docking nine homology-modeled targets. Each panel ranks models with an interface RMSD in Å between 0 Å and 10 Å. Each model is colored by CAPRI rank,^63^ with incorrect models in gray, acceptable models in blue, medium-quality models in orange, and high-quality models in red. The bound refined native models are shown in purple. The CXCR4 chemokine receptor (3OE0) which was docked unsuccessfully is shown in (d) as a membrane view and (e) as a top view. The crystal structure is shown in gray and the best-scoring docked model is shown in blue.

Fig. 8c shows the performance of the nine homology-model docking targets. Each panel plots all 5,000 models by interface RMSD and interface score, with each model colored by CAPRI criteria. In four of the nine cases (1E12, 1M56, 1ZOY, and 3RVY), the energy landscape has a funnel pattern, with incorrect models scoring high and near-native models receiving the lowest score. Thus, *franklin2019* recognized near-native interfaces. Further, scores for the refined native structures (purple) are near the bottom of the funnel of sampled docked structures. The five best-scoring models are shown in Fig. S9 and S10.

For the five remaining homology-modeled targets, RosettaMPDock fails to recognize the correct interface. For two cases (3CHX and 3OE0), the docking program did not sample any near-native models. Amongst the remaining cases (1M0L, 2QJY, and 3KLY), the energy function prefers acceptable or incorrect models over near-native models. One example of a challenging case is the CXCR chemokine receptor (Fig. 8d-e, 3OE0), where the low-scoring models have an incorrect bilayer orientation.

### Summary of *franklin2019* successes and challenges

Above, we described and demonstrated protocols for twelve scientific benchmark tests. To integrate the test results, we established a threshold or optimization goal for the summary metrics (Table 3).

**Table 3:** Summary of current benchmark test performance criteria

| # | Test | Metrics | Exp. Error | Threshold/Goal |
| --- | --- | --- | --- | --- |
| 1 | Tilt angle | tilt | 5° | 10° |
| 2 | Rotation angle | rotation | 12° | 12° |
| 3 | Protein orientation | tilt/depth | 10°/2.5Å | 10°/2.5Å |
| 4 | Hydrophobic thickness | thickness | 2.5 Å | 2.5 Å |
| 5 | $\Delta G_{w,l}$ at constant pH | $\Delta G_{ins}$ | 1.4 kcal/mol | max $R^2$ |
| 6 | $\Delta\Delta G_{w,l}$ with pH shift | $\Delta\Delta G_{pH-ins}$ | 0.1 kcal/mol | max $R^2$ |
| 7 | $\Delta\Delta G^{mut}$ | $\Delta\Delta G_{mut}$ | 0.6 kcal/mol | max $R^2$ |
| 8 | Sequence recovery | $R_s/KL$ | NA | max $R_s$/min KL |
| 9 | Side chain distribution | AUΔC | NA | min AUΔC |
| 10 | Decoy discrimination | NA | $D$ | 5 Å |
| 11 | Helix kinks | kink | NA | 10° |
| 12 | Docking | $\langle N_5 \rangle / \langle E_{1\%} \rangle$ | NA | 3/2 |

Together, the benchmark results reveal strengths and pitfalls of a current energy function. For instance, test #1 (Fig. 1) showed effective prediction of single-transmembrane helix tilt angles. In addition, test #9 (Fig. 5) showed the energy function can predict many bilayer depth-dependent amino acid preferences. In contrast, test #5 (Fig. 3) illuminated the need to capture shifts in proton avidity for titratable sites in the low dielectric bilayer, and test #4 (Fig. 2) suggested the need for better implicit membrane interfacial representations to accurately predict hydrophobic thickness. Test #11 (Fig. 2) highlighted pitfalls in the hydrogen bonding model. Further, tests #4 and #5 probed the balance of forces in the overall energy function. Together, the results suggest areas for future energy function optimization.

## Discussion

We developed 12 scientific benchmarks to evaluate energy functions for membrane protein modeling and design. Our approach addresses the challenge of limited experimental data by bundling many small sets to achieve a large and feature-rich dataset. The tests account for effects of the heterogeneous lipid bilayer on protein orientation, stability, sequence, and structure. The tests encompass wide-ranging modeling tasks from ΔΔ*G* calculations to protein-protein docking and design. As a step forward from single-test validation^7–9,12^ or four-test validation,^16,18^ we anticipate that these benchmarks will accelerate development of the next generation of membrane protein energy functions.

An interesting consequence of the membrane is the type of data that can be used for benchmarking. For soluble proteins, energy functions for molecular modeling and force fields for molecular dynamics rely on a combination of small molecule thermodynamic data and known macromolecular structures. For membrane proteins, all-atom force fields also use physical chemistry data to derive parameters for different lipid types. However, the analogous small molecule data are difficult to obtain for implicit membrane simulations because organic solvents do not sufficiently mimic the properties of heterogeneous biological membranes.^67^ Further, it is challenging to rigorously measure thermodynamic properties in lipid bilayers. Hydrophobic length and membrane protein re-orientation can be observed on short timescales so it is accessible via molecular dynamics.^9^ However, molecular dynamics is computationally expensive for modeling long-timescale biological phenomena such as some conformational changes and protein binding. By using a Monte Carlo approach with an implicit biologically-realistic membrane, we enabled a faster calculation of both thermodynamic and structural properties. For a single run, the tests require no more than 10,000 CPU hours. As a result, it is practical to iteratively run the tests to maintain reproducibility and for continuous optimization. For this purpose, the tests run continuously on the Rosetta Benchmark Server (https://benchmark.graylab.jhu.edu/) and the source code is publicly distributed through GitHub (https://github.com/rfalford12/Implicit-Membrane-Energy-Function-Benchmark) to make the tests accessible to all membrane protein modeling developers. We hope these resources will help the community share standardized metrics for evaluating membrane protein energy functions.

This first multi-faceted benchmark is a base upon which the quantity and quality of test data can be extended. For quantity, improvements in structure determination will increase the number and diversity of known structures to benefit both sequence and structure tests. In contrast, the data are still sparse for stability and orientation since these values are not revealed when structures are determined in detergents. The paucity of data limits splitting the data into a training and a testing set, a key practice for demonstrating generalizability of models. On the other hand, quality is determined by various factors including the resolution of crystal and NMR structures, the uncertainty of stability measurements, and the rigor of assumptions made to obtain and analyze the data. The challenge of quality and quantity is well illustrated by considering the datasets of ΔΔG of mutation measurements. Kroncke *et al.*^7^ compiled a large dataset of ΔΔG of mutation measurements. However, the reference state for each measurement varied (e.g., lipid composition, ion concentration in aqueous phase), making it challenging to compare. As a result, we used a smaller set of ΔΔG measurements to improve quality. Comparing measurements is a consistent challenge of resolving the complexity of membrane proteins performed in different lipid compositions and environments.

As much as possible, our tests evaluate energy functions with experimental reference data. By transitivity, the energy function can only be as accurate as these reference data. For soluble proteins, the wealth of biophysical and structural information compensates. As of April 2020, there are more than 160,000 structures deposited in the Protein Databank.^68^ Consequently, soluble protein energy functions can be validated on large and diverse datasets.^21,69^ In contrast, only 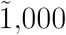 membrane protein structures are deposited in the Protein Data Bank.^5^ A central challenge is that many membrane proteins are not naturally abundant and cannot be reconstituted into a membrane mimetic. Thus, substantial improvements to the energy function will require both reliable benchmarks and significant advances in experimental methods.

Important future work includes developing a framework to perform a global optimization of the energy function. A possible approach is to develop a series of objective functions that define the relationship between the threshold and performance for all targets in the dataset. For example, an objective function to define the performance of the tilt angle test (Test #1) could be formulated as

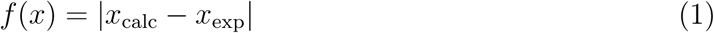

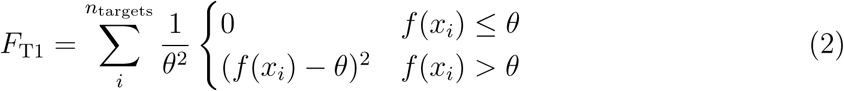

where *x*_calc_ is the predicted tilt angle, *x*_exp_ is the experimentally measured tilt angle, *i* is an iterator over all targets in the dataset, and *θ* = 10° is an allowable amount of error. Objective functions for the other 11 tests could be similarly constructed, and a weighted sum could serve as a loss function to train on the whole benchmark set. Deep learning has recently piqued the interest of the structural biology community.^70^ Currently, deep learning approaches require large, high-quality datasets. Recently, Wang *et al.*^71^ used transfer learning to develop a transmembrane protein structure prediction algorithm that relies on a soluble protein contact prediction algorithm^72^ and the Deep CNF transmembrane topology prediction algorithm.^73^ Our benchmark data can be used with approaches like these, particularly toward incorporating the all-important lipid composition and specificity features to move toward more accurate and biologically realistic implicit energy functions.

## Author Information

### Conflicts

Dr. Gray is an unpaid board member of the Rosetta Commons. Under institutional participation agreements between the University of Washington, acting on behalf of the Rosetta Commons, Johns Hopkins University may be entitled to a portion of revenue received on licensing Rosetta software including programs described here. As a member of the Scientific Advisory Board, JJG has a financial interest in Cyrus Biotechnology. Cyrus Biotechnology distributes the Rosetta software, which may include methods described in this paper.

### Data and Software Availability

All benchmark protocols are available within the Rosetta Software suite at https://www.rosettacommons.org to all non-commercial users for free and to commercial users for a fee. The benchmark datasets are available through the Rosetta Benchmark Server (distributed) and the Membrane Energy Function Benchmark Project GitHub Repository at https://github.com/rfalford12/Implicit-Membrane\-Energy-Function-Benchmark. Current benchmark performance is available through the Rosetta Scientific Benchmark server at https://benchmark.graylab.jhu.edu/revisions?branch=scientific.

## Acknowledgement

This research was supported by a Hertz Foundation Fellowship (R.F.A.), a National Science Foundation Graduate Research Fellowship (R.F.A.) and NIH Grant GM-072281 (R.F.A. and J.J.G.). Computational resources were provided by the Maryland Advanced Research Computing Center (MARCC) and the Texas Advanced Computing Center (TACC). We wish to thank Karen Fleming and Pat Fleming for helpful discussions and feedback on the manuscript. We also wish to thank Julia Koehler Leman and Sergey Lyskov for development of the Rosetta benchmark server.

## Supporting Information Available

Detailed methods and supplementary figures are available online

