## Supplementary Information for "Diverse scientific benchmarks for implicit membrane energy functions"

### Diverse scientific benchmarks reveal optimization imperatives for implicit membrane energy functions

#### SUPPLEMENTARY METHODS

**Benchmark curation** 2

**Protocols** 3

Test #1-3: Membrane protein orientation . . . 3
Test # 4: Hydrophobic length . . . 3
Test # 5: $\Delta\Delta G$ of peptide insertion at constant pH . . . 4
Test #6: $\Delta\Delta G$ of peptide insertion upon pH shift . . . 4
Test #7: $\Delta\Delta G$ of mutation . . . 4
Test #8-9: Sequence features . . . 4
Test #10: Native structure discrimination . . . 5
Test # 11: Helix Kinks . . . 5
Test #12: Protein-protein docking . . . 6

#### LIST OF FIGURES

S1 Tilt angle prediction for biological peptides with a single transmembrane domain. . . . 8
S2 Tilt angle prediction for designed peptides with a single transmembrane domain. . . . 8
S3 Rotation angle prediction for surface-adsorbed biological peptides. . . . 9
S4 Change in energy with different bilayer thickness for hydrophobic length estimate. . . . 10
S5 Comparison between experimentally measured $\Delta\Delta G^{mut}$ and *franklin2019* water-to-bilayer score. . . . 10
S6 Contributions of component energies to the $\Delta\Delta G^{mut}$ for mutations to tryptophan. . . . 11
S7 Contributions of component energies to the $\Delta\Delta G^{mut}$ for mutations to tyrosine. . . . 12
S8 Difference in the depth-dependent side chain distribution between native and designed sequences. . . . 13
S9 Comparison of five lowest scoring docked membrane protein complexes from membrane view. . . . 14
S10 Comparison of five lowest scoring docked membrane protein complexes from top view. . . . 15

#### BENCHMARK CURATION

For each benchmark set, all experimental X-Ray and NMR structures were downloaded from the Orientations of Proteins in Membranes database(1). Each PDB coordinate file was cleaned to remove any non-protein atoms, excluding essential ligands where noted. Experimental structures were not available for designed helical peptides. In these instances, we generated structural models by threading the sequences onto an ideal helix scaffold. We defined the transmembrane topology for each protein structure as follows. For proteins with a single transmembrane segment, the topology was defined as a single span beginning at the N-terminus and ending at the C-terminus. For proteins with multiple transmembrane segments, the topology was calculated from the structure using the `mp_span_from_pdb` application(2) with a membrane thickness of 30 Å. The benchmark sets are listed in Supplementary Tables summarized in Table S1.

**Table S1:** Targets for membrane protein energy function benchmarks

| Description | # of Targets | Listing | Refs. |
|---|---|---|---|
| **Tilt angle measurements** | | | |
| Transmembrane helical peptides | 5 | Table S1 | (3) |
| Surface-adsorbed helical peptides | 8 | Table S2 | (3) |
| Transmembrane helical peptides | 3 | Table S3 | (4) |
| Surface-adsorbed helical peptides | 1 | Table S4 | (5) |
| **Orientation and Hydrophobic Length** | | | |
| Multipass $\alpha$ and $\beta$ proteins w/ known orientation | 34 | Table S5 | (6) |
| **Stability measurements** | | | |
| Poly leucine peptides | 5 | Table S9 | (7) |
| pH-sensitive peptides | 16 | Table S10 | (8) |
| Structures of OmpLA and PagP | 2 | Table S6 | (9–11) |
| **High-resolution structure prediction** | | | |
| $\alpha$ proteins with multiple states | 4 | Table S11 | (12–15) |
| Single-pass homo-dimers | 18 | Table S12 | (16) |
| Multi-pass homo- and hetero-dimers | 48 | Table S13 | (17) |
| Multi-pass high-resolution $\alpha$-helical structures | 5 | Table S15 | (18, 19) |
| Sequentially-distant $\alpha$ and $\beta$ proteins | 129 | Table S17 | (20) |

### PROTOCOLS

#### Test #1-3: Membrane protein orientation

To predict depth and tilt angle, the `MembraneOrientationSampler` application was used to exhaustively sample and score possible orientations. The orientation sampler performs an initial side chain repacking and energy minimization along all torsion angles ($\psi$,$\phi$ and $\chi$). The pose is transformed to an initial orientation at 0° and -60 Å from the membrane center (80 Å for larger proteins). Then, all orientations are sampled and scored between depths of -60 Å to 60 Å with a 1 Å step size and tilt angles between 0-360° with a 1° step size. For transmembrane peptides, the rotation is relative to the $z$-axis; whereas, for surface-adsorbed peptides, the rotation is relative to the $x$-axis, as triggered by the "interface" option. The `MembraneOrientationSampler` protocol was implemented with the following command:

```
Rosetta/main/source/bin/rosetta_scripts.macosclangrelease
   -in:file:s 1a11.pdb                          # Input PDB File
   -mp:setup:spanfiles single_TM_mode           # Spanning topology
   -mp:lipids:composition DLPC                  # Lipid composition
   -parser:protocol orientation_sampler.xml     # XML Script protocol
   -parser:script_vars                          # Energy function to use
        sfxn_weights=franklin2019
```

The contents of orientation_sampler.xml are listed below.

```
<ROSETTASCRIPTS>
   <SCOREFXNS>
      <ScoreFunction name="s" weights="%%sfxn_weights%%"/>
   </SCOREFXNS>
   <TASKOPERATIONS>
      <RestrictToRepacking name="rtrp"/>
      <ExtraRotamersGeneric name="extra_chi" ex1="1" ex2="1" extrachi_cutoff="0"/>
   </TASKOPERATIONS>
   <MOVERS>
      <AddMembraneMover name="add_memb" />
      <TransformIntoMembraneMover name="transform_into_memb" />
      <PackRotamersMover name="pack_rotamers" scorefxn="s" task_operations="rtrp"/>
      <MinMover name="minimize_struc" scorefxn="s" chi="1" bb="1" jump="0" />
      <MembraneEnergyLandscapeSampler name="landscape_test" scorefxn="s" interface="0" />
   </MOVERS>
   <PROTOCOLS>
      <Add mover_name="add_memb"/>
      <Add mover_name="transform_into_memb"/>
      <Add mover_name="pack_rotamers"/>
      <Add mover_name="minimize_struc"/>
      <Add mover_name="landscape_test"/>
   </PROTOCOLS>
</ROSETTASCRIPTS>
```

The application outputs a data file containing all of the orientations and corresponding energies. The orientation corresponding to the lowest energy is considered the most favored orientation.

#### Test # 4: Hydrophobic length

For each target, the protein was positioned at the membrane orientation predicted by the Orientations of Proteins in Membranes database(1). An initial round of side-chain repacking and energy minimization was performed to remove steric clashes. Then, the protein was scored at all hydrophobic lengths ranging between 0-40 Åwith a step size of 1 Å. This protocol was implemented in the `find_optimal_hydrophobic_thk` application and run using the command below.

```
Rosetta/main/source/bin/find_optimal_hydrophobic_thk.macosclangrelease
   -in:file:s 2QJY.pdb                          # Input PDB File
```

```
   -mp:setup:spanfiles from_structure         # Spanning topology
   -mp:lipids:composition DLPC                # Lipid composition
   -score:weights franklin2019                # Energy function to use
   -pack_missing_sidechains 0                 # Do not pack before membrane is setup
```

##### Test # 5: $\Delta\Delta G$ of peptide insertion at constant pH

The $\Delta\Delta G$ of insertion was computed as the difference between the peptide energy when spanning the bilayer $\Delta G_{\text{bilayer}}$ and the energy of the unfolded peptide in solution $\Delta G_{\text{water}}$. To compute $\Delta G_{\text{bilayer}}$ and $\Delta G_{\text{water}}$, the MembraneOrientationSampler was used to exhaustively sample and score all possible orientations (see Test #1-3). Then, the $\Delta G_{\text{bilayer}}$ was taken to be the lowest energy orientation among all orientations with a depth between ±15 Å. $\Delta G_{\text{water}}$ was taken to be the lowest energy orientation among all orientations with a depth greater or less than ±15 Å.

##### Test #6: $\Delta\Delta G$ of peptide insertion upon pH shift

First, we generated a mapping between orientations and energies at both pH values. We used the same protocol as Tests # 1-3 in combination with the pH energy framework (21). To adapt the protocol for pH, we added the `e_pH` score term to the *franklin2019* weights file with a weight of 1.0. In addition, we added two flags to the command line: `-pH_mode true -value_pH <4 or 8>`. Then, the $\Delta\Delta G_{\text{ins-pH}}$ was computed as the difference between the peptide energy when spanning the membrane at pH 4 and the peptide energy when in solution at pH 8.

##### Test #7: $\Delta\Delta G$ of mutation

To predict the $\Delta\Delta G$ of mutation in the membrane environment, we used the Rosetta MPddG application. The protocol is a fixed backbone $\Delta\Delta G$ prediction protocol, similar to Kellogg *et al.* that optimizes all side chain conformations within 8 Åof the mutated residue. Then, the $\Delta\Delta G$ was computed as the difference in Rosetta Energy Units (REU) between the mutant and native conformation. The $\Delta\Delta G_{\text{mut}}$ was computed using the `predict_ddG_of_mutation.py` script using the command below.

```
python3 predict_ddG_of_mutation.py
   --in_pdb 1QD6.pdb                                # Input PDB File
   --spanfile from_structure                        # Spanning topology
   --mutation_list OmpLA_Moon_Fleming_set.dat       # List of mutations to evaluate
   --energy_fxn franklin2019                        # Energy function to use
```

##### Test #8-9: Sequence features

Low energy sequences were identified through fixed-backbone protein redesign using the `RosettaFixBB` application. The `RosettaFixBB` protocol is a rotamer and sequence optimization protocol that samples rotamers for all positions and then seeks the lowest-energy assignment of rotamers to the structure using a Monte Carlo simulated annealing protocol (22). The `RosettaFixBB` protocol was implemented using the command line below.

```
Rosetta/main/source/bin/fixbb.macosclangrelease
   -in:file:s 1AFO.pdb                      # Input pdb file
   -mp:setup:spanfiles from_structure         # Spanning topology
   -mp:lipids:composition DLPC                # Lipid composition
   -score:weights franklin2019                # Energy function to use
   -out:file:scorefile 1AFO.sc              # Path to output score file
   -in:membrnae                   # Initialize the membrane framework
   -pack_missing_sidechains 0                 # Do not pack before membrane setup
   -nstruct 1                       # Number of models to generate
```

#### Test #10: Native structure discrimination

The Rosetta `MPRelax` protocol (2) is adds the context of the membrane to the `FastRelax` protocol. `FastRelax` is an algorithm that refines the protein structure through five cycles of side chain packing and energy minimization in torsion space ($\phi$, $\psi$, and $\omega$). For each subsequent cycle, the repulsive energy is allowed to increase from 0.2 to a full weight of 1.0. `FastRelax` is adapted for the membrane in two ways. First, the energy function includes energy terms that account for the lipid bilayer. Second, minimization is also performed on the degrees of freedom that describe the orientation of the protein in the bilayer. The `MPRelax` protocol was implemented with the command below.

```
Rosetta/main/source/bin/rosetta_scripts.macosclangrelease
   -in:file:s BRD7.pdb                          # Input PDB File
   -mp:setup:spanfiles from_structure           # Spanning topology
   -mp:lipids:composition DLPC                  # Lipid composition
   -mp:lipids:has_pore false                    # For targets without pores
   -nstruct 5                                   # Number of models to generate
   -relax:constrain_relax_to_start_coords       # Keep the backbone intact
   -out:file:scorefile BRD7.sc                  # Path to output scorefile
   -parser:protocol mp_relax.xml                # XML Script protocol
   -parser:script_vars                          # Energy function to use
        sfxn_weights=franklin2019
```

The contents of `mp_relax.xml` are given below.

```
<ROSETTASCRIPTS>
   <SCOREFXNS>
      <ScoreFunction name="memb_hires" weights="%%sfxn_weights%%" />
   </SCOREFXNS>
   <MOVERS>
      <AddMembraneMover name="add_memb"/>
      <MembranePositionFromTopologyMover name="init_pos"/>
      <FastRelax name="fast_relax" scorefxn="memb_hires" repeats="8"/>
   </MOVERS>
   <PROTOCOLS>
      <Add mover="add_memb"/>
      <Add mover="init_pos"/>
      <Add mover="fast_relax"/>
   </PROTOCOLS>
   <OUTPUT scorefxn="memb_hires" />
</ROSETTASCRIPTS>
```

#### Test # 11: Helix Kinks

To capture backbone hinge motions, a conformational ensemble was generated using normal mode analysis (NMA) (23). NMA provides information about accessible equilibrium modes, assuming the system is stabilized by harmonic potentials. In Rosetta, NMA is implemented as the `NormalModeRelaxMover`. First, the protocol mixes motion along the first five normal modes with a 1 Åperturbation. To prevent non-physical bond lengths and angles, a scoring term was added to penalize deviations from ideal bond lengths and angles. This step iterated with the relax protocol described in *Native structure discrimination*. We accessed this through the Rosetta XML interface, called `RosettaScripts`. The command line is given below.

```
Rosetta/main/source/bin/rosetta_scripts.macosclangrelease
   -in:file:s 3FV7.pdb                 # Input PDB File
   -nstruct 100                        # Number of models to generate
   -out:file:scorefile 3FV7.sc         # Path to output scorefile
   -parser:protocol nma.xml            # XML Script protocol
```

The contents of `nma.xml` are listed below.

```
<ROSETTASCRIPTS>
   <SCOREFXNS>
      <ScoreFunction name="bn15_cart" weights="ref2015_cart" />
   </SCOREFXNS>
   <MOVERS>
      <NormalModeRelax name="nma" cartesian="true" centroid="false"
      scorefxn="bn15_cart" nmodes="5" mix_modes="true" pertscale="1.0"
      randomselect="false" relaxmode="relax" nsample="20"
      cartesian_minimize="false" />
   <PROTOCOLS>
      <Add mover="nma" />
   </PROTOCOLS>
   <OUTPUT scorefxn="bn15_cart" />
</ROSETTASCRIPTS>
```

##### Test #12: Protein-protein docking

Prior to docking, the side chains of each partner were repacked in isolation. Here, $X_1$ is the chain ID for the receptor and $X_2$ is the chain ID for the receptor. The pre-packing step was performed with the command line below.

```
Rosetta/main/source/bin/docking_prepack_protocol.macosclangrelease
   -in:file:s 1AFO.pdb                          # Input PDB File
   -mp:setup:spanfiles from_structure           # Spanning topology
   -mp:lipids:composition DLPC                  # Lipid composition
   -mp:lipids:has_pore false                    # For targets without pores
   -score:weights franklin2019                  # Energy function to use
   -out:file:scorefile 1AFO_ppk.sc              # Path to output score file
   -docking:partners X1_X2                      # Chains for the receptor and ligand
   -pack_missing_sidechains 0                   # Do not pack before membrane setup
   -nstruct 1                                   # Number of models to generate
```

Next, the pre-packed structure was used as the starting structure for local, rigid-body docking. We used `MPDock` which adapts `RosettaDock` version 3.0 for docking within the membrane by performing translations and rotations in consideration of the plane of the bilayer. The docking protocol was performed with the command line below.

```
Rosetta/main/source/bin/mpdocking.macosclangrelease
   -in:file:s 1AFO_ppk.pdb                      # Input prepacked pdb file
   -in:file:native 1AFO_native.pdb              # Input native bound conformation
   -mp:setup:spanfiles from_structure           # Spanning topology
   -mp:lipids:composition DLPC                  # Lipid composition
   -mp:lipids:has_pore false                    # For targets without pores
   -score:weights franklin2019                  # Energy function to use
   -out:file:scorefile 1AFO_ppk.sc              # Path to output score file
   -docking:partners X1_X2                      # Chains for the receptor and ligand
   -docking:dock_pert 3 8                       # Magnitude of translation and rotation
   -pack_missing_sidechains 0                   # Do not pack before membrane setup
   -nstruct 5000                                # Number of models to generate
```

In addition, to evaluate the quality of docked models the bound states were refined using the command line below.

```
Rosetta/main/source/bin/mpdocking.macosclangrelease
   -in:file:s 1AFO_ppk.pdb                      # Input prepacked pdb file
   -in:file:native 1AFO_native.pdb              # Input native bound conformation
   -mp:setup:spanfiles from_structure           # Spanning topology
   -mp:lipids:composition DLPC                  # Lipid composition
   -mp:lipids:has_pore false                    # For targets without pores
```

```
-score:weights franklin2019              # Energy function to use
-out:file:scorefile 1AFO_ppk.sc          # Path to output score file
-docking:partners X1_X2                  # Chains for the receptor and ligand
-docking:dock_pert 3 8                   # Magnitude of translation and rotation
-docking_local_refine                    # Only perform local refinement
-pack_missing_sidechains 0               # Do not pack before membrane setup
-nstruct 5000                            # Number of models to generate
```

#### SUPPLEMENTAL FIGURES

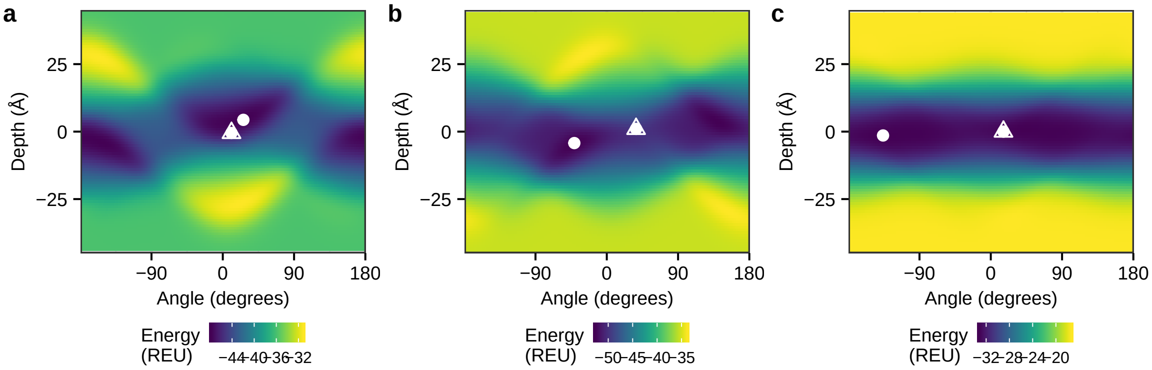

**Figure S1:** Tilt angle prediction for biological peptides with a single transmembrane domain. A mapping between all sampled orientations and energies for (a) Acetylcholine receptor segment M2, (b) Influenza A segment M2, and (c) VPU-forming domain of HIV-1 protein. Here, tilt angle relative to the membrane normal (in degrees) is on the $x$-axis and depth relative to the membrane center is on the $y$-axis. Each grid point is colored by *franklin2019* energy, with low energies colored in dark blue and high energies colored in yellow. Further, each grid point represents a 1Å and 1°increment. the lowest energy predicted orientation is shown as a white circle, and where applicable the experimentally measured orientation is shown as a white triangle.

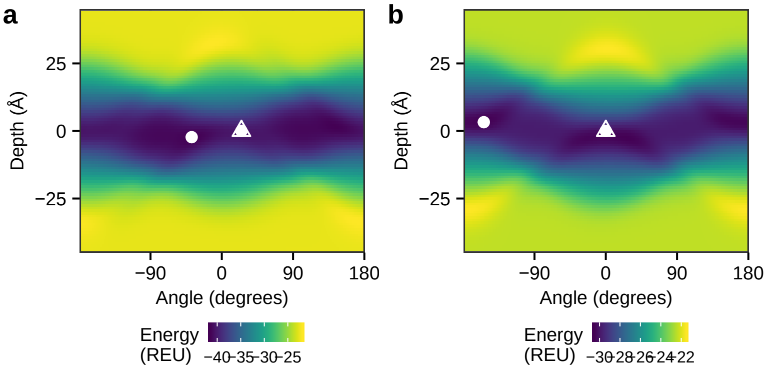

**Figure S2:** Tilt angle prediction for designed peptides with a single transmembrane domain. A mapping between all sampled orientations and energies for (a) WALP23 and (b) a poly-alanine helix with flanking tyrosine residues. Here, tilt angle relative to the membrane normal (in degrees) is on the $x$-axis and depth relative to the membrane center is on the $y$-axis. Each grid point is colored by *franklin2019* energy, with low energies colored in dark blue and high energies colored in yellow. Further, each grid point represents a 1Å and 1°increment. the lowest energy predicted orientation is shown as a white circle, and where applicable the experimentally measured orientation is shown as a white triangle.

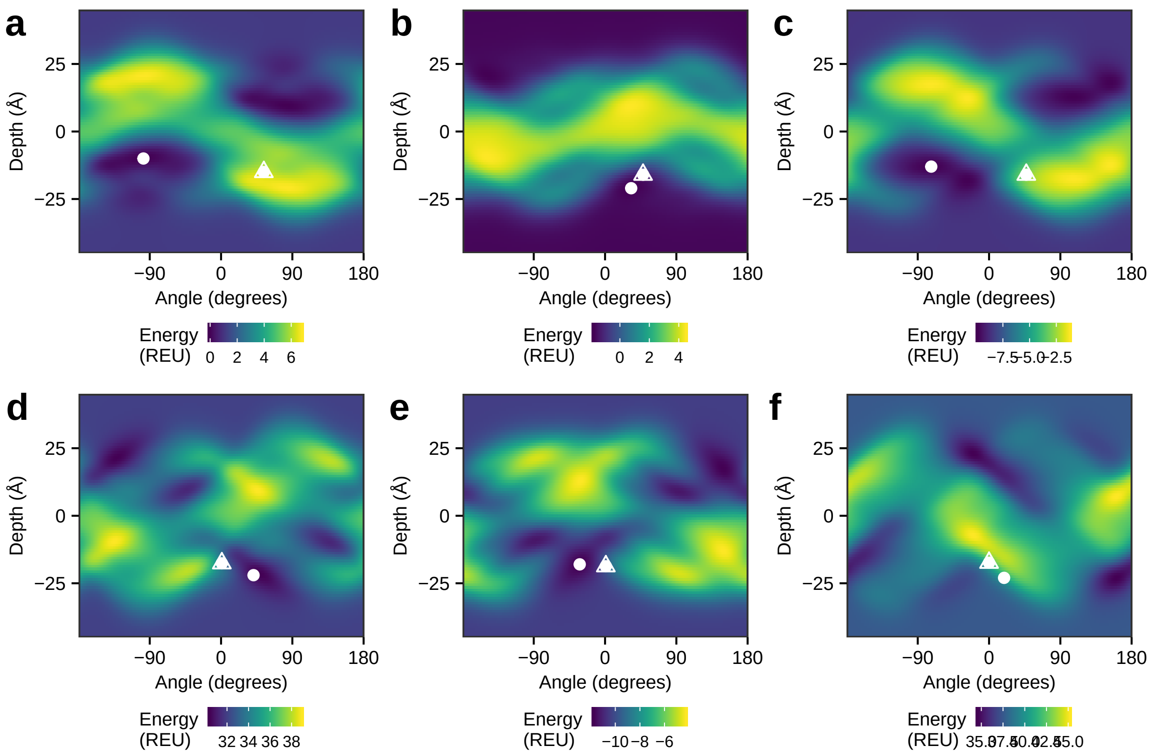

**Figure S3:** Rotation angle prediction for surface-adsorbed biological peptides. A mapping between all sampled orientations and energies for (a) Ovispirin 1 (b) Novispirin G10, (c) Novispirin T2, (d) Magainin-cercopin hybrid, (e) Magainin-cercopin hyrbrid P2, and (f) Magainin-cercopin hybrid P1. Here, rotation angle relative to the membrane surface plane (in degrees) is on the $x$-axis and depth relative to the membrane center is on the $y$-axis. Each grid point is colored by *franklin2019* energy, with low energies colored in dark blue and high energies colored in yellow. Further, each grid point represents a 1Å and 1°increment. the lowest energy predicted orientation is shown as a white circle, and where applicable the experimentally measured orientation is shown as a white triangle.

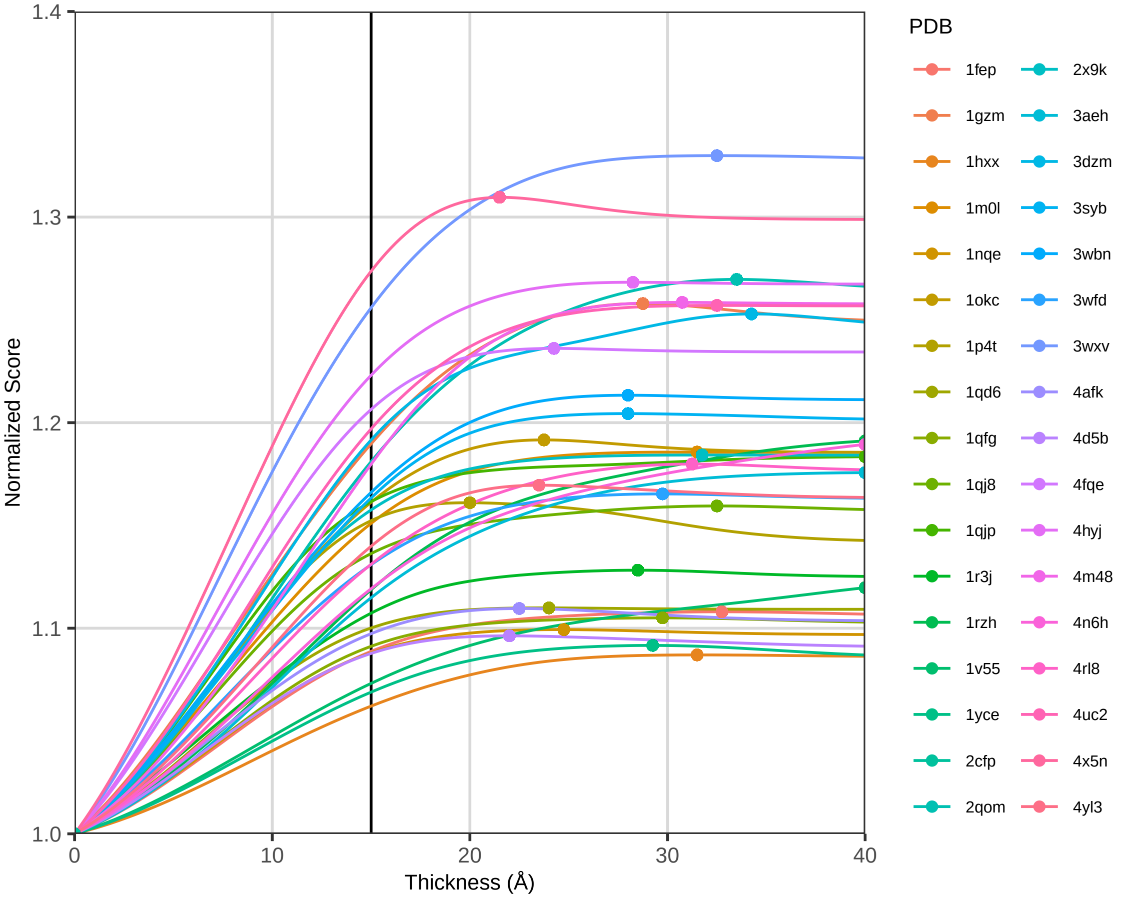

**Figure S4:** Change in energy with different bilayer thickness for hydrophobic length estimate. The energy is normalized relative to the maximum score for the protein. Each curve represents the change in energy with bilayer thickness for a different target. Each point represents the minimum energy bilayer thickness.

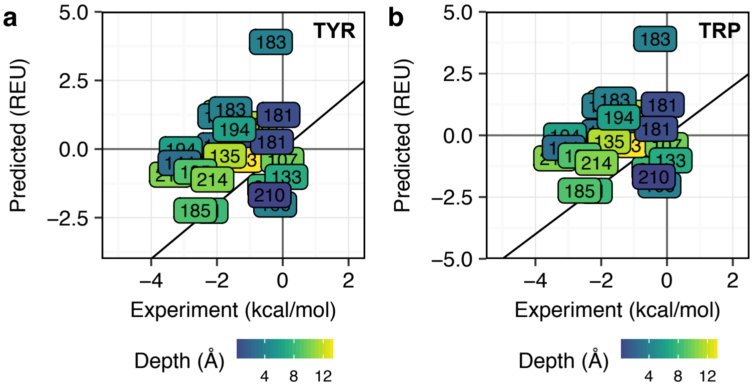

**Figure S5:** Comparison between experimentally measured $\Delta\Delta G^{\mathrm{mut}}$ and *franklin2019* water-to-bilayer score. Values for mutations to tyrosine and tryptophan are shown in (a) and (b) respectively. Each position is colored by depth relative to the membrane center plane in Å on a scale from blue (closer to the center) to yellow (closer to the interface/water barrier). The $y = x$ line is shown as a bold black line. The experimentally measured values were taken from (10).

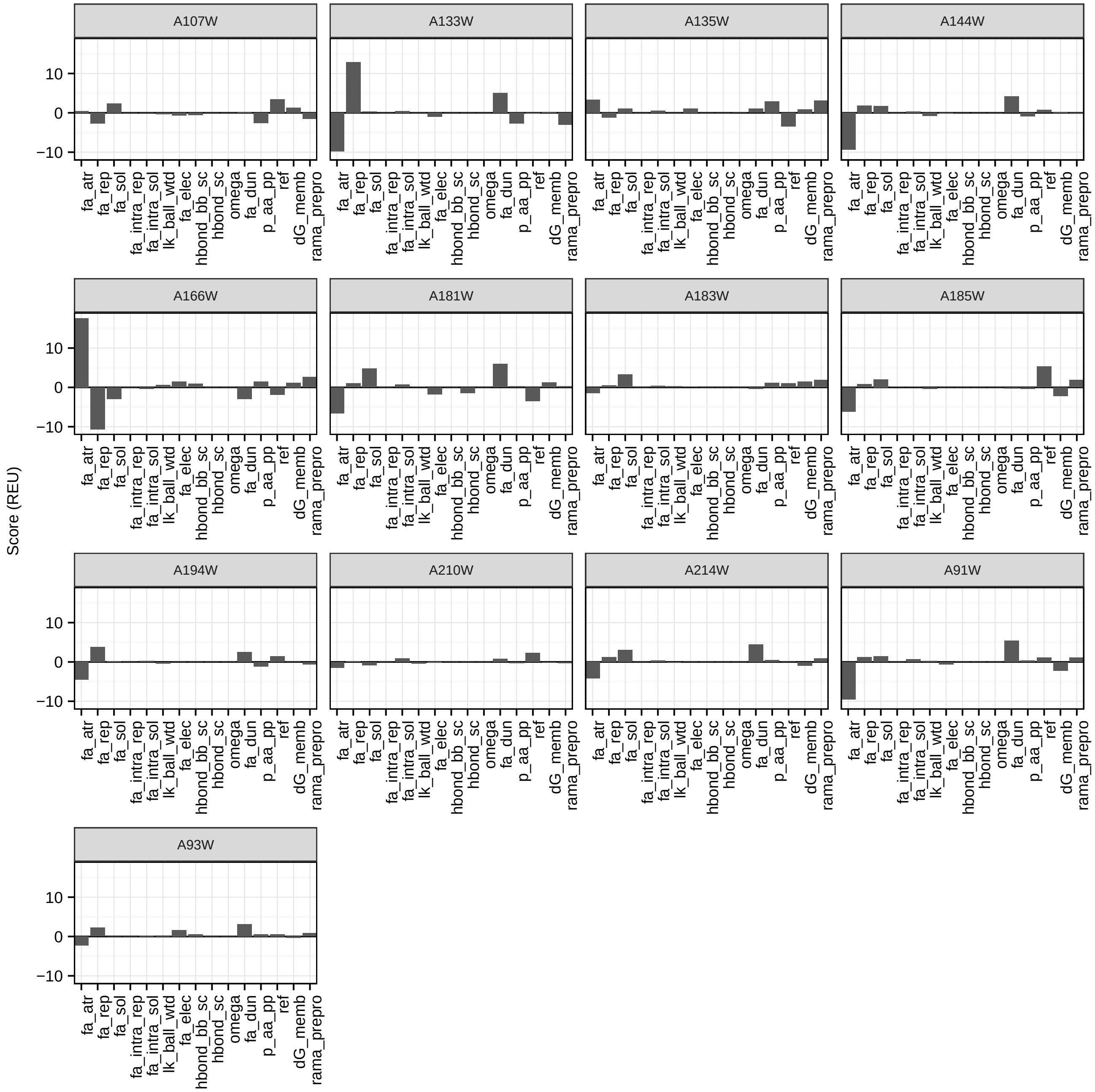

**Figure S6:** Contributions of component energies to the $\Delta\Delta G^{\mathrm{mut}}$ for mutations to tryptophan. Energies are only listed for contributions > 0.1 REU to the $\Delta\Delta G_{\mathrm{mut}}$. These energy terms include the van der Waals attractive energy (fa_atr), repulsive energy (fa_rep), Lazaridis-Karplus solvation energy (fa_sol), intra-residue repulsive energy (fa_intra_rep), intra-residue Lazaridis-Karplus solvation energy (fa_intra_sol), orientation-dependent component of Lazaridis-Karplus solvation energy (lk_ball_wtd), Coulomb electrostatics energy (fa_elec), backbone-side chain hydrogen bonding energy (hbond_bb_sc), side chain to side chain hydrogen bonding energy (hbond_sc), omega torsion energy (omega), Dunbrack rotamer energy (fa_dun), knowledge-based backbone $\phi, \psi$ energy (rama_prepro, p_aa_pp), amino acid reference energy (ref), and *franklin2019* water-to-bilayer energy (dG_memb).

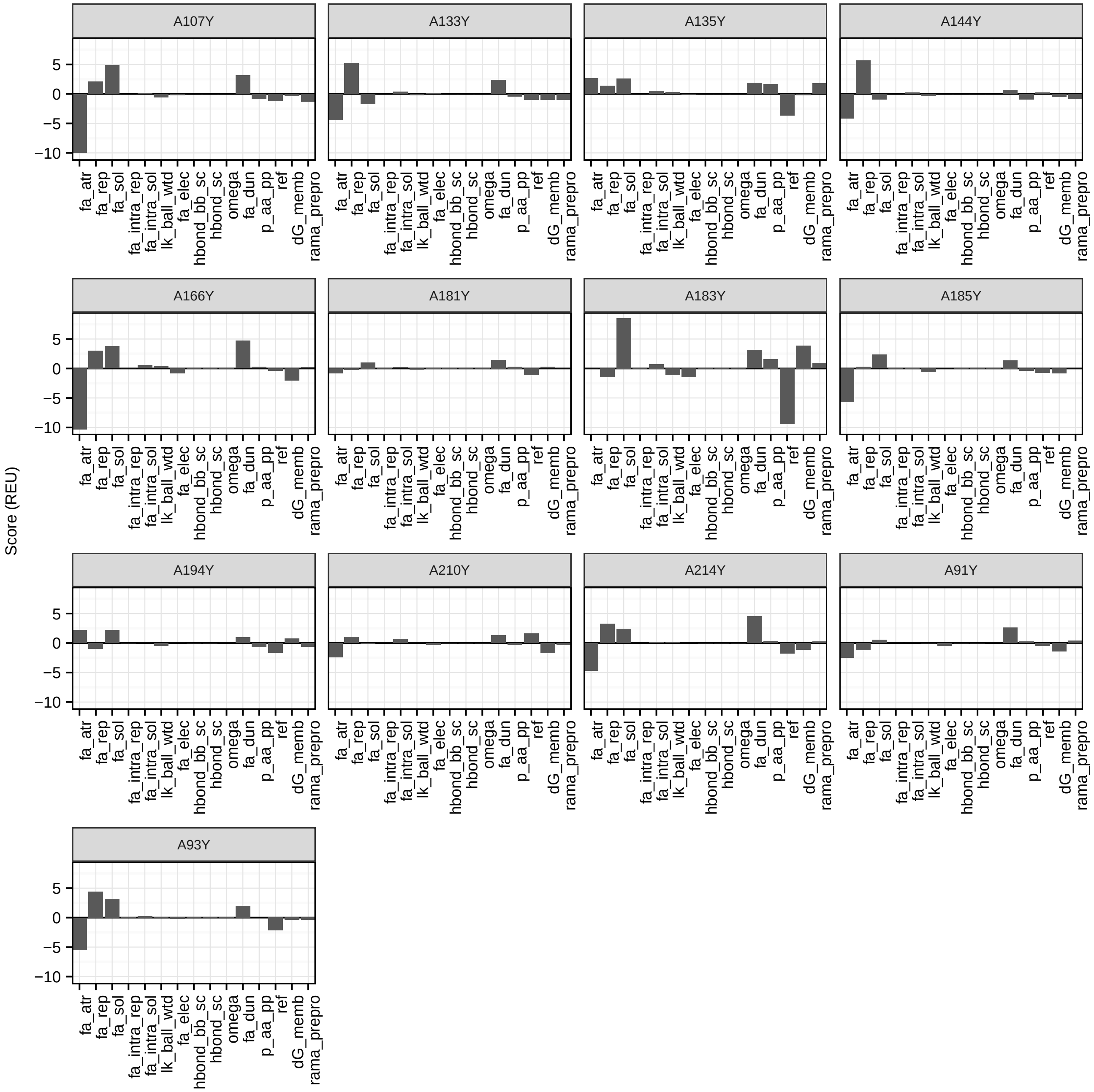

**Figure S7:** Contributions of component energies to the $\Delta\Delta G^{mut}$ for mutations to tyrosine. Energies are only listed for contributions > 0.1 REU to the $\Delta\Delta G_{mut}$. These energy terms include the van der Waals attractive energy (fa_atr), repulsive energy (fa_rep), Lazaridis-Karplus solvation energy (fa_sol), intra-residue repulsive energy (fa_intra_rep), intra-residue Lazaridis-Karplus solvation energy (fa_intra_sol), orientation-dependent component of Lazaridis-Karplus solvation energy (lk_ball_wtd), Coulomb electrostatics energy (fa_elec), backbone-side chain hydrogen bonding energy (hbond_bb_sc), side chain to side chain hydrogen bonding energy (hbond_sc), omega torsion energy (omega), Dunbrack rotamer energy (fa_dun), knowledge-based backbone $\phi, \psi$ energy (rama_prepro, p_aa_pp), amino acid reference energy (ref), and *franklin2019* water-to-bilayer energy (dG_memb).

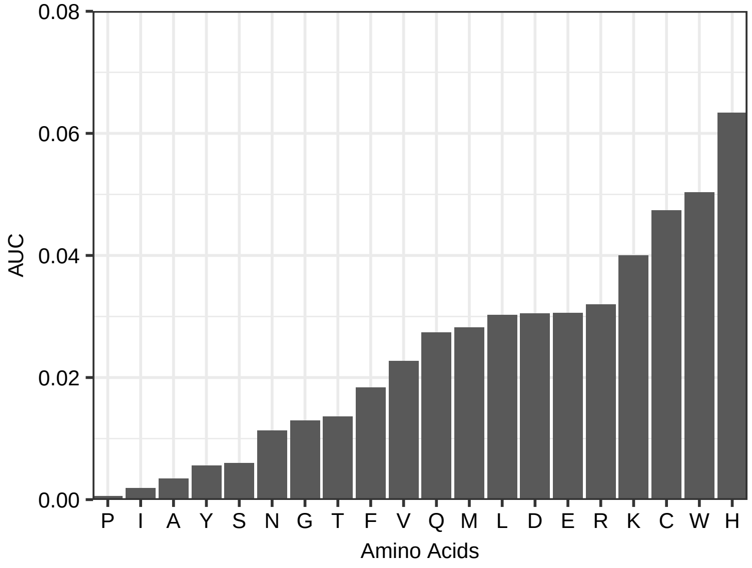

**Figure S8:** Difference in the depth-dependent side chain distribution between native and designed sequences. The difference in distributions is calculated as the difference between the area-under-the-curve for the native and designed amino acid distributions. The area under the curve was computed using a trapezoidal rule.

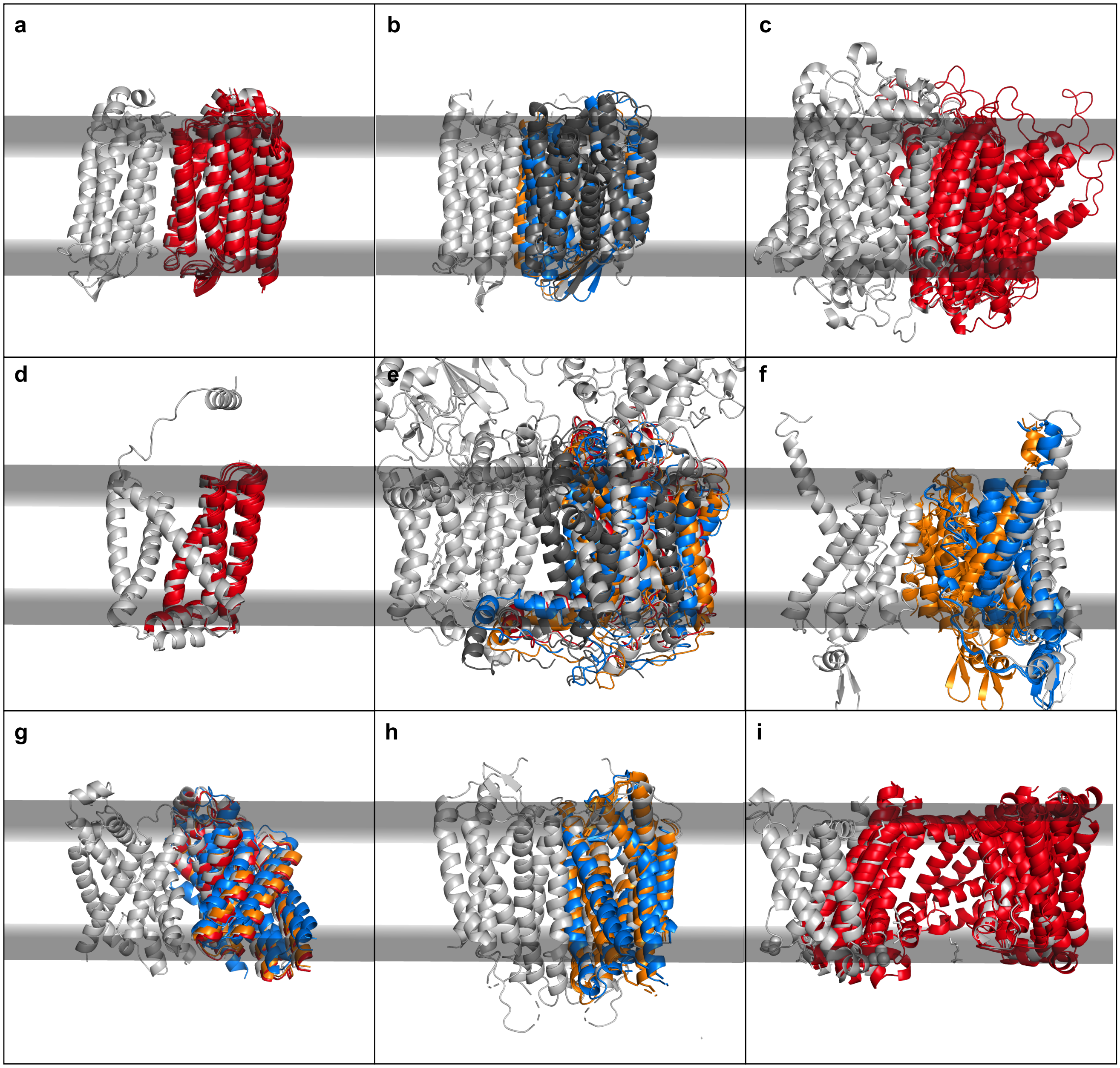

**Figure S9:** Comparison between the native and five lowest scoring docked membrane protein complexes from membrane view. The native membrane protein complex is colored in gray and the docked models are colored by CAPRI criteria. Here, high-quality models are in red, medium-quality models are in orange, acceptable models are in blue, and incorrect models are in dark grey. Each panel shows docked models for one of the nine unbound docking targets: (a) halorhodopsin, (b) bacteriorhodopsin, (c) cytochrome C oxidase, (d) mitochondrial respiratory complex II, (e) cytochrome bc1, (f) methane monooxygenase, (g) pentameric formate channel, (h) CXCR4 chemokine receptor, (i) NavAb voltage-gated sodium channel.

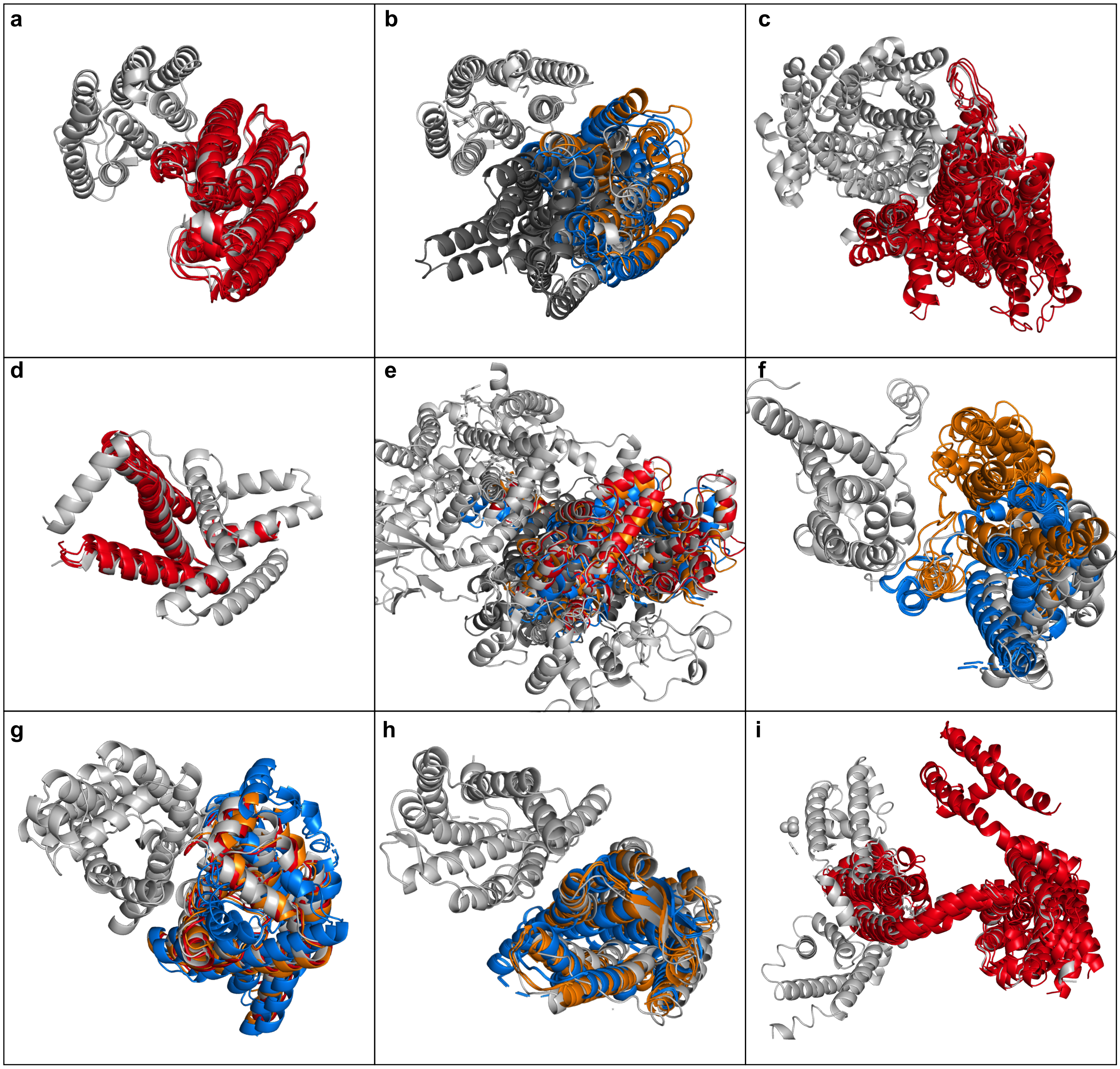

**Figure S10:** Comparison between the native and five lowest scoring docked membrane protein complexes from top view. The native membrane protein complex is colored in gray and the docked models are colored by CAPRI criteria. Here, high-quality models are in red, medium-quality models are in orange, acceptable models are in blue, and incorrect models are in dark grey. Each panel shows docked models for one of the nine unbound docking targets: (a) halorhodopsin, (b) bacteriorhodopsin, (c) cytochrome C oxidase, (d) mitochondrial respiratory complex II, (e) cytochrome bc1, (f) methane monooxygenase, (g) pentameric formate channel, (h) CXCR4 chemokine receptor, (i) NavAb voltage-gated sodium channel.

#### SUPPLEMENTAL TABLES

**Table S2:** Transmembrane peptide targets for tilt angle test

| Name | PDB | Membrane | Sequence | Tilt Angle(°) | Ref. |
|---|---|---|---|---|---|
| Acetylcholine M2 | 1a11 | DPC | GSEKMSTAISVLLAQAVFLLLTSQR | 11 | (24) |
| Influenza A M2 | 1mp6 | DMPC | SSDPLVVAASIIGILHLILWILDRL | 38 ± 3 | (25) |
| VPU domain | 1pje | DOPC:DOPG | MQPIQIAIVALVVAIIIAIVVWSIVIIEGRGGKKKK | 16 | (26) |
| NMDA Receptor | 2nr1 | DPC | GSNGDALTLSSAMWFSWGVLLNSGIGE | 40 ± 2 | (24) |
| WALP23 | – | DOPC | GWWLALALALALALALALALWWA | 24.8 | (27) |
| PolyA-Y | – | – | AAAYAAAAAAAAAAAAAAAYA | – | – |
| PolyA-W | – | – | AAAWAAAAAAAAAAAAAAAAWA | – | – |

**Table S3:** Membrane surface-adsorbed peptide targets for rotation angle test

| Name | PDB | Membrane | Sequence | Tilt Angle (°) | Ref. |
|---|---|---|---|---|---|
| Magainin | 2mag | DPC | `GIGKFLHSAKKFGKAFVGEIMNSX` | 87 ± 7 | (28) |
| Ovispirin 1 | 1hu5 | 2,2,2-trifluoroethanol | `KNLRRIIRKIIHIIKKYG` | 94 ± 11 | (29) |
| Novispirin G10 | 1hu6 | 2,2,2-trifluoroethanol | `KNLRRIIRKGIHIIKKYG` | 88 ± 6 | (29) |
| Novispirin T2 | 1hu7 | 2,2,2-trifluoroethanol | `KNLRRITRKIIHIIKKYG` | 87 ± 12 | (29) |
| Magainin-cercopin hybrid | 1f0e | DPC micelles | `KWKLFKKIPKFLHSAKKFX` | 91 ± 8 | (30) |
| Magainin-cercopin hybrid P2 | 1f0f | DPC micelles | `KWKLFKKIKFLHSAKKFX` | 88 ± 4 | (30) |
| Magainin-cercopin hybrid P1 | 1f0g | DPC micelles | `KLKLFKKIGIGKFLHSAKKFX` | 90 ± 10 | (30) |
| Leucine-lysine repeat | – | – | `LKKLLKLLKKLLKLLKKLLKLLKKL` | – | (5) |

**Table S4:** Multi-pass $\alpha$-helical and $\beta$-barrel proteins for orientation and hydrophobic thickness test

| PDB | Protein | Class | Localization |
|---|---|---|---|
| 1fep | Ferric enterobactin receptor | $\beta$ | *E. coli* Gram-negative outer membrane |
| 1gzm | Bovine rhodopsin | $\alpha$ | Eukaryotic plasma membrane |
| 1hxx | OmpF porin channel | $\beta$ | *E. coli* Gram-negative outer membrane |
| 1m0l | Bacteriorhodopsin | $\alpha$ | Archea membrane |
| 1nqe | Outer membrane cobalambin transporter BtuB | $\beta$ | *E. coli* Gram-negative outer membrane |
| 1okc | Mitochondrial ATP/ADP Carrier | $\alpha$ | Mitochondria inner membrane |
| 1p4t | Neisserial surface protein A | $\beta$ | *E. coli* Gram-negative outer membrane |
| 1qd6 | Outer membrane protein phospholipase A | $\beta$ | *E. coli* Gram-negative outer membrane |
| 1qfg | Ferric hydroxamate receptor (FhuA) | $\beta$ | *E. coli* Gram-negative outer membrane |
| 1qj8 | Outer membrane protein OmpX | $\beta$ | *E. coli* Gram-negative outer membrane |
| 1qjp | Outer membrane protein A (OmpA) | $\beta$ | *E. coli* Gram-negative outer membrane |
| 1r3j | Potassium channek KcsA | $\alpha$ | *E. coli*Gram-negative outer membrane |
| 1rzh | Photosynthetic reaction center | $\alpha$ | *E. coli* Gram-negative outer membrane |
| 1v55 | Cytochrome C oxidase | $\alpha$ | Mitochondria inner membrane |
| 1yce | Rotor ring of F-type Na+-ATP-ase | $\alpha$ | *E. coli* Gram-negative inner membrane |
| 2cfp | Sugar-free lactose permease | $\alpha$ | *E. coli* Gram-negative inner membrane |
| 2qom | EspP autotransporter beta-domain | $\beta$ | *E. coli* Gram-negative outer membrane |
| 2x9k | Outer membrane protein G (OmpG) | $\beta$ | *E. coli* Gram-negative outer membrane |
| 3aeh | Autotransporter Hbp | $\beta$ | *E. coli* Gram-negative outer membrane |
| 3dzm | Outer membrane protein TtoA | $\beta$ | *E. coli* Gram-negative outer membrane |
| 3syb | Outer membrane carboxylate channel (OpdP) | $\beta$ | *E. coli* Gram-negative outer membrane |
| 3wbn | MATE multidrug transporter | $\alpha$ | Archea membrane |
| 3wfd | Nitric oxide reductase | $\alpha$ | *E. coli*Gram-negative outer membrane |
| 3wxv | Adopinectin receptor 1 | $\alpha$ | Eukaryotic plasma membrane |
| 4afk | Alginate transporter AlgE | $\beta$ | *E. coli* Gram-negative outer membrane |
| 4d5b | Outer membrane protein CymA | $\beta$ | *E. coli* Gram-negative outer membrane |
| 4fqe | Oligogalacturonate-specific KdgM porin | $\beta$ | *E. coli* Gram-negative outer membrane |
| 4hyj | Proteorhodopsin | $\alpha$ | *E. coli* Gram-negative outer membrane |
| 4m48 | Dopamine transporter | $\alpha$ | Eukaryotic plasma membrane |
| 4n6h | Delta opioid receptor | $\alpha$ | Eukaryotic plasma membrane |
| 4rl8 | COG4313 outer membrane channel | $\beta$ | *E. coli* Gram-negative outer membrane |
| 4uc2 | TSPO transporter protein | $\alpha$ | *E. coli* Gram-negative inner membrane |
| 4x5n | SemiSWEET transporter | $\alpha$ | *E. coli* Gram-negative inner membrane |
| 4yl3 | mPGES-1 inhibitor complex | $\alpha$ | Endoplasmic reticulum membrane |

**Table S5:** Reference OPM values for hydrophobic thickness, tilt angle, and depth

| PDB | Protein | Length (Å) | Angle (°) | Depth (Å) |
|---|---|---|---|---|
| 1fep | Ferric enterobactin receptor | 24.3 | 1 | 12.8 |
| 1gzm | Bovine rhodopsin | 32.2 | 11 | -0.4 |
| 1hxx | OmpF porin channel | 24 | 0 | -6.6 |
| 1m0l | Bacteriorhodopsin | 31.8 | 0 | -1.7 |
| 1nqe | Outer membrane cobalambin transporter BtuB | 23.4 | 5 | 9.8 |
| 1okc | Mitochondrial ATP/ADP Carrier | 29.5 | 14.1 | -4.4 |
| 1p4t | Neisserial surface protein A | 24.9 | 22 | 6.5 |
| 1qd6 | Outer membrane protein phospholipase A | 23.9 | 0 | -5.2 |
| 1qfg | Ferric hydroxamate receptor (FhuA) | 24.7 | 5 | -12.4 |
| 1qj8 | Outer membrane protein OmpX | 23.6 | 12 | -7.3 |
| 1qjp | Outer membrane protein A (OmpA) | 25.4 | 11 | -3.8 |
| 1r3j | Potassium channek KcsA | 34.8 | 0 | 2.1 |
| 1rzh | Photosynthetic reaction center | 31.8 | 2 | 8.6 |
| 1v55 | Cytochrome C oxidase | 28 | 0 | 2.5 |
| 1yce | Rotor ring of F-type Na+-ATP-ase | 37 | 0 | -4.2 |
| 2cfp | Sugar-free lactose permease | 31.1 | 2.2 | -4.4 |
| 2qom | EspP autotransporter beta-domain | 25.1 | 6 | 7.7 |
| 2x9k | Outer membrane protein G (OmpG) | 24.7 | 5 | -7.6 |
| 3aeh | Autotransporter Hbp | 25.2 | 4 | -7.6 |
| 3dzm | Outer membrane protein TtoA | 28.5 | 16 | 11.4 |
| 3syb | Outer membrane carboxylate channel (OpdP) | 23.6 | 8 | -6.8 |
| 3wbn | MATE multidrug transporter | 31.8 | 8 | -0.7 |
| 3wfd | Nitric oxide reductase | 31.7 | 11.6 | -0.4 |
| 3wxv | Adopinectin receptor 1 | 32.8 | 15 | -2 |
| 4afk | Alginate transporter AlgE | 24.8 | 3 | 7 |
| 4d5b | Outer membrane protein CymA | 23.5 | 8.3 | 10.6 |
| 4fqe | Oligogalacturonate-specific KdgM porin | 22.2 | 2 | 3.3 |
| 4hyj | Proteorhodopsin | 30 | 15 | -1.5 |
| 4m48 | Dopamine transporter | 30.8 | 12 | 1.9 |
| 4n6h | Delta opioid receptor | 34 | 14 | -9.1 |
| 4rl8 | COG4313 outer membrane channel | 23.4 | 6 | -6.2 |
| 4uc2 | TSPO transporter protein | 30.4 | 1 | 4.6 |
| 4x5n | SemiSWEET transporter | 36.8 | 1 | 0.8 |
| 4yl3 | mPGES-1 inhibitor complex | 29.8 | 0 | -3.9 |

**Table S6:** Sequences and measured insertion energies for designed poly-leucine peptides

| Name | Sequence | Membrane | $\Delta G_{\text{insert}}$ |
|---|---|---|---|
| GL5 | GLLLLLRLLLLLG | POPC | 2.1 |
| GL6 | GLLLLLLRLLLLLLG | POPC | 0.5 |
| GL7 | GLLLLLLLRLLLLLLLG | POPC | -0.5 |
| GL8 | GLLLLLLLLRLLLLLLLLG | POPC | -1.5 |
| GWL6 | GWLLLLLLRLLLLLLLG | POPC | -0.5 |

**Table S7:** Sequences and measured insertion energies for designed pH-sensitivie peptides

| Name | Sequence | $\Delta G_{\text{insert}}$ |
|---|---|---|
| v1 | ACEDQNPYWARYADWLFTTPLLLLDLALLVDG | 2.17 |
| v2 | ACEDQNPYWRAYADLFTPLTLLDLLALWDG | 0.28 |
| v3 | ACDDQNPWRAYLDLLFPTDTLLLDLLW | 2.23 |
| v4 | ACEEQNPWRAYLELLFPTETLLLELLW | 1.72 |
| v5 | ACDDQNPWARYLDWLFPTDTLLLDL | 2.31 |
| v6 | CDNNNPWRAYLDLLFPTDTLLLDW | 1.93 |
| v7 | ACEEQNPWARYLEWLFPTETLLLEL | 2.39 |
| v8 | CEEQQPWAQYLELLFPTETLLLEW | 2.19 |
| v9 | CEEQQPWRAYLELLFPTETLLLEW | 2.07 |
| v10 | ACEDQNPWARYADWLFPTTLLLLD | 1.79 |
| v11 | ACEEQNPWARYAEWLFPTTLLLLE | 1.94 |
| v12 | ACEDQNPWARYADLLFPTTLAW | 1.95 |
| v13 | ACEEQNPWARYAELLFPTTLAW | 1.15 |
| v14 | TEDADVLLALDLLLLPTTFLWDAYRAWYPNQECA | 1.78 |
| v15 | CDDDDDNPNYWARYANWLFTTPLLLLNGALLVEAEET | 0.82 |
| v16 | CDDDDDNPNYWARYAPWLFTTPLLLLPGALLVEAEET | 1.47 |

**Table S8:** Targets for helix kink prediction

| Target | Localization | State | PDB |
|---|---|---|---|
| Potassium channel KcsA | Bacterial gram-positive plasma membrane | closed | 1r3j |
| | Bacterial gram-positive plasma membrane | open-inactive | 3f7v |
| B2-adrenergic GPCR | Eukaryotic plasma membrane | inactive | 2rh1 |
| | Eukaryotic plasma membrane | active | 3p0g |
| Adiponectin receptor 1 | Eukaryotic plasma membrane | closed | 3wxv |
| | Eukaryotic plasma membrane | open | 5lxg |
| Platelet activating receptor | Eukaryotic plasma membrane | closed | 5zkp |
| | Eukaryotic plasma membrane | open | 5zkq |
